# A domain-swapped CaMKII conformation facilitates linker-mediated allosteric regulation

**DOI:** 10.1101/2024.03.24.586494

**Authors:** Bao V. Nguyen, Can Özden, Kairong Dong, Oguz Can Koc, Ana P. Torres-Ocampo, Noelle Dziedzic, Daniel Flaherty, Jian Huang, Saketh Sankara, Nikki Lyn Abromson, Diana R. Tomchick, Rafael A Fissore, Jianhan Chen, Scott C Garman, Margaret M Stratton

## Abstract

Memory formation, fertilization, and cardiac function rely on precise Ca^2+^ signaling and subsequent Ca^2+^/calmodulin-dependent protein kinase II (CaMKII) activation. Ca^2+^ sensitivity of the four CaMKII paralogs in mammals has been linked to the length of the variable linker region that is a hot spot for alternative splicing. In this study, we determined that the position of charged residues within the linker modulates the Ca^2+^/CaM sensitivity. We present an X-ray crystal structure of the full-length CaMKIIδ holoenzyme consisting of domain-swapped dimers within a dodecameric complex. In this structure, the kinase domain of one subunit is docked onto the hub domain of an adjacent subunit, providing an additional interface within the holoenzyme. Mutations at the hub equatorial and lateral interfaces led to alterations in the stoichiometry of CaMKII holoenzyme as well as Ca^2+^/CaM sensitivity. Using molecular dynamics (MD) to compare domain-swapped to non-domain-swapped CaMKIIs, we demonstrate that the domain-swapped conformation facilitates an interaction between the calmodulin binding region and the linker region. Based on MD simulations and small-angle X-ray scattering (SAXS) measurements, we propose a model where the position of charges on the linker region drives an interaction with the regulatory segment that modulates the degree of autoinhibition. Finally, we use live-cell imaging to show that the activation profiles we observe in vitro are recapitulated in cells. Our findings provide a new framework for understanding allosteric regulation of CaMKII by the linker region in Ca^2+^-sensitive cells.

## INTRODUCTION

Ca^2+^/calmodulin-dependent protein kinase II (CaMKII) is critical for signaling during long-term memory formation, fertilization, and cardiac pacemaking ^1–3^. There are four CaMKII genes (α, β, δ, γ) in mammals that are expressed differentially throughout the body, where CaMKIIα and β are predominantly neuronal, CaMKIIγ is expressed in muscle tissue, oocytes and sperm, and CaMKIIδ is enriched in cardiomyocytes. All CaMKIIs share the same domain organization: an N-terminal kinase domain, followed by a regulatory segment and variable linker region, and finally the C-terminal hub (oligomerization) domain. CaMKII is activated upon an increase in intracellular Ca^2+^ sensed by calmodulin (CaM); Ca^2+^/CaM competitively binds the CaMKII regulatory segment thereby making the substrate binding site available ^4^. The hub domain forms multiple stoichiometries of CaMKII holoenzymes: dodecameric, tetradecameric and hexadecameric complexes ^5–11^.

CaMKII regulation and Ca^2+^/CaM sensitivity have been linked to neurologic dysfunction and dilated cardiomyopathy as evidenced by *de novo* point mutations identified in the catalytic and hub domains of CaMKII ^12–15^. Mutations in the kinase domain and regulatory segment perturb Ca^2+^/CaM sensitivity as well as stability in some cases. One of these mutations, H477Y, is located in the hub domain and was identified in a patient with neurodevelopmental syndrome ^16^. This hub mutation resulted in oligomer dissociation coupled with reduced neuronal activity, which is consistent with previous work connecting the hub domain identity to activity regulation ^10^.

The kinase and the hub domains are connected by a variable linker region. The four human CaMKII paralogs have many splice sites in the linker region, creating upwards of 70 variants expressed in the human hippocampus ^10^. The linker sequence and length are variable, from zero (*i.e*., no linker) to 217 residues, affording them different biophysical properties and post-translational modifications ^17,18^. Much focus has been placed on understanding the functional implications of the length aspect of the linker region. Previous studies have shown that the linker length modulates Ca^2+^/CaM sensitivity of CaMKIIα, but not CaMKIIβ ^6,10,19^. The linker region of CaMKIIα is encoded by exon 14, exon 15, and exon 18. In human hippocampus, CaMKIIα with a 30-residue linker (CaMKIIα-30) comprised of exon 14 and exon 18 is the predominant splice variant, while CaMKIIα variants containing no linker or all three exons exist to a lower level ^10^. CaMKIIβ has the longest possible linker of 217 residues comprised of exons 13, 14, 16, 17, 18, 19, 20, 21, although it is undetectable in human hippocampus and is as sensitive to Ca^2+^/CaM as CaMKIIβ-no linker ^10^. Even though emerging data support the notion that alternative splicing of CaMKII is a regulated process ^20,21^, the implications of sequence variation within the linker region on CaMKII structure and activity are unclear. While many structures of CaMKII kinase and hub domains as isolated domains have been determined ^22,23^, they do not visualize the linker region, which is predicted to be disordered. The first crystal structure of the intact CaMKIIα-0 revealed a new conformation (deemed compact state) where the kinase domain was docked onto the hub domain of the same subunit ^6^. This crystal structure provided a clear explanation for biochemical observations that CaMKIIα-0 (no linker) requires a high concentration of Ca^2+^/CaM for activation.

Small-angle X-ray scattering (SAXS) and cryo-electron microscopy (cryo-EM) have been employed to assess the structure of CaMKIIα with a 30-residue linker which activates more easily, at lower concentrations of Ca^2+^/CaM. SAXS showed a significant difference between CaMKIIα-0 and CaMKIIα-30, where CaMKIIα-30 had a larger envelope, indicating that the kinases were farther from the hub instead of docked as seen in CaMKIIα-0 ^6^. Cryo-EM showed that subunits within a CaMKIIα-30 complex did adopt a compact conformation, but not all subunits as seen in the CaMKIIα-0 structure ^10^. These results imply that extending the linker length increases flexibility, but still allows an equilibrium between compact and extended states.

These biochemical and structural data have led to a model for CaMKIIα where longer linkers increase kinase flexibility farther from the hub domain (extended conformation), increasing accessibility to Ca^2+^/CaM ^6,8^. Shorter linkers (*i.e*., no linker) reduce kinase flexibility leading to a stronger interaction between the kinase and hub domains (compact conformation), decreasing accessibility to Ca^2+^/CaM. Here, we further examined whether the sequence of the linker region also impacts Ca^2+^/CaM sensitivity. We employed a combination of biochemistry, SAXS, X-ray crystallography, molecular dynamics (MD) simulations, and live-cell CaMKII activity measurements to frame our biochemical results in light of CaMKII conformation and regulation. Herein, we present the structure of an intact CaMKIIδ-0 holoenzyme, which reveals an unambiguous domain-swapped dimer conformation. In the context of this new structure combined with activity measurements and MD simulations, we propose a model for how the linker sequence, composition, and orientation due to domain swapping affect the accessibility of the Ca^2+^/CaM binding site to dictate CaMKII sensitivity to calcium.

## RESULTS

### The variable linker sequence affects Ca^2+^/CaM sensitivity

We hypothesized that the composition of the linker region, in addition to its length, could directly modulate the sensitivity of CaMKII to Ca^2+^/CaM. To test this, we changed the linker sequence in CaMKIIα but maintained similar lengths and compared this to other variants: CaMKIIβ and CaMKIIδ (both 95% similar to CaMKIIα). All CaMKII variants encode exons 14 and 18 in the genetic background. CaMKIIα has the shortest encoded variable linker region, which only includes exons 14, 15, and 18 ^10^ (**Fig. 1A**). Exons 14 and 18 are similar lengths, comprised of 14 and 16 residues, respectively, but have different sequences (**Fig. 1B**). We constructed variants of CaMKIIα, β, and δ containing only exon 14 or only exon 18 as the linker region from the corresponding genetic sequence (i.e. CaMKIIα with α exon 14, etc.). For all variants, inclusion of exon 14 significantly increased the Ca^2+^/CaM sensitivity compared to variants containing exon 18. CaMKII variants containing just exon 14 had EC_50_ values ∼20 nM whereas variants containing just exon 18 had EC_50_ values ranging from 91 – 193 nM (**Fig. 1C-E**). CaMKIIα displayed the largest shift in EC_50_ value, from 17 nM to 193 nM, so we performed further experiments using CaMKIIα.

**Figure 1.**
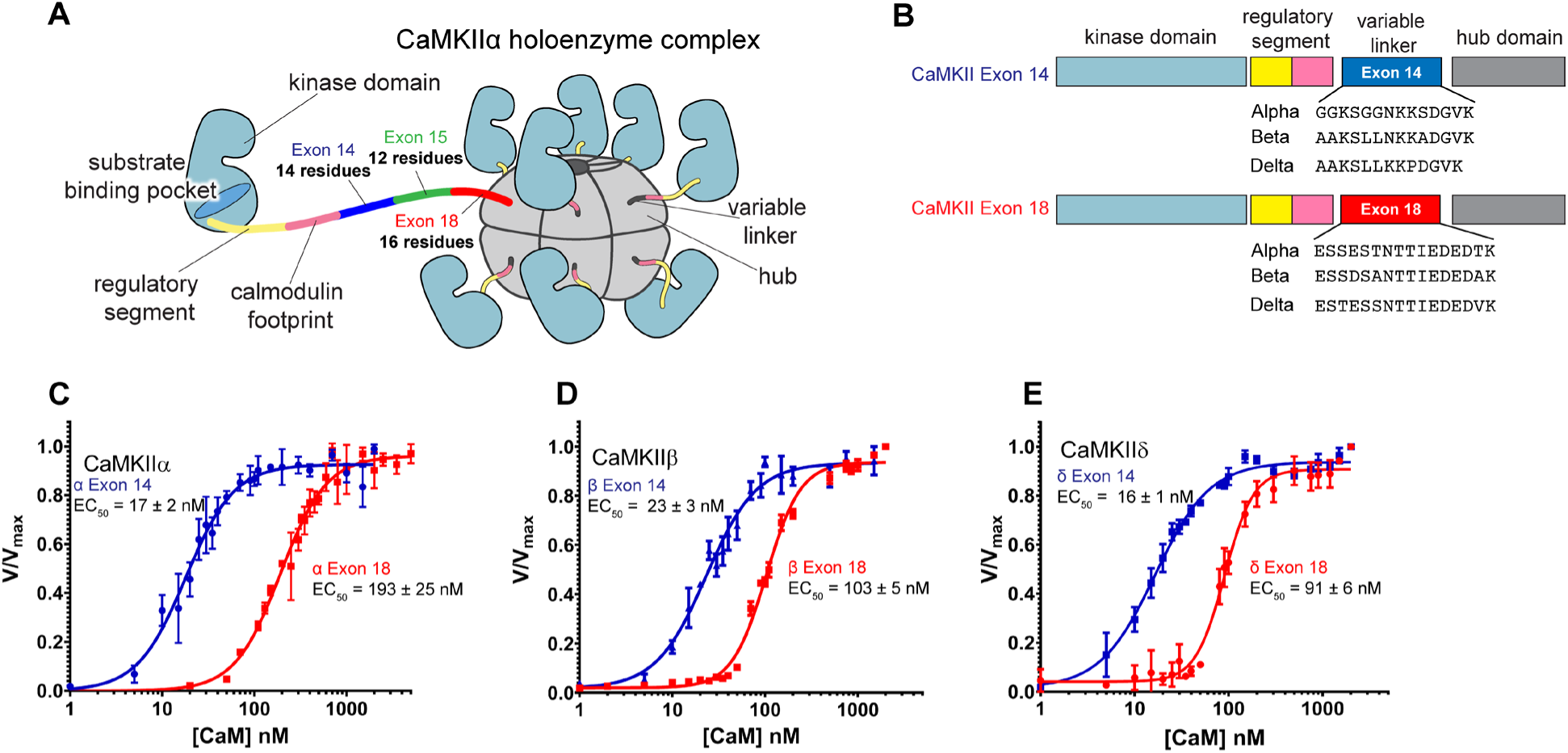
More than the length of the variable linker region affects Ca^2+^/CaM sensitivity. (A) Overview of CaMKIIα holoenzyme organization containing the linker region that is made of exon 14, exon 15, and exon 18. (B) Subunit architecture and sequences of exon 14 and exon 18 in the variable linker region of CaMKIIα, CaMKIIβ, and CaMKIIδ. (C-E) Kinase assays of CaMKIIα (C), CaMKIIβ (D), and CaMKIIδ (E) comparing the sensitivity to Ca^2+^/CaM between CaMKII with exon 14 and with exon 18 as the linker region. Each kinase assay comparison was done on the same day. Each data point represents the mean ± SD, n=3.

### Phosphorylation of the variable linker does not account for difference in Ca^2+^/CaM sensitivity

There are two striking differences in the sequences of exons 14 and 18. One is the number of potential autophosphorylation sites, and the second is the net charge (**Fig. 2A**). We first tested whether autophosphorylation occurs under our in vitro kinase assay by performing phosphoproteomic analysis of CaMKIIα upon incubation with ATP/Mg^2+^ and Ca^2+^/CaM for 1 minute at room temperature. Peptides were analyzed using LC/MS/MS and showed that all Ser and Thr residues on exon 18 were autophosphorylated under these conditions *in vitro* (**Fig. S1A-F**). We created a CaMKII mutant with all Ser and Thr residues in exon 18 mutated to Ala (S/TΔA mutant). If phosphorylation played a role in causing the increased sensitivity, we would expect this mutant to be left-shifted compared to WT exon 18. Instead, this non-phosphorylatable mutant required more Ca^2+^/CaM for activation (EC_50_ = 350 ± 25 nM) compared to WT exon 18 (**Fig. S1G**). From these data, we concluded that although these residues can be phosphorylated *in vitro*, this is not why inclusion of exon 18 makes it more difficult for this variant to bind Ca^2+^/CaM. For the subsequent experiments, we used the S/TΔA mutant to remove the confounding variable of autophosphorylation of the linker unless stated otherwise.

**Figure 2.**
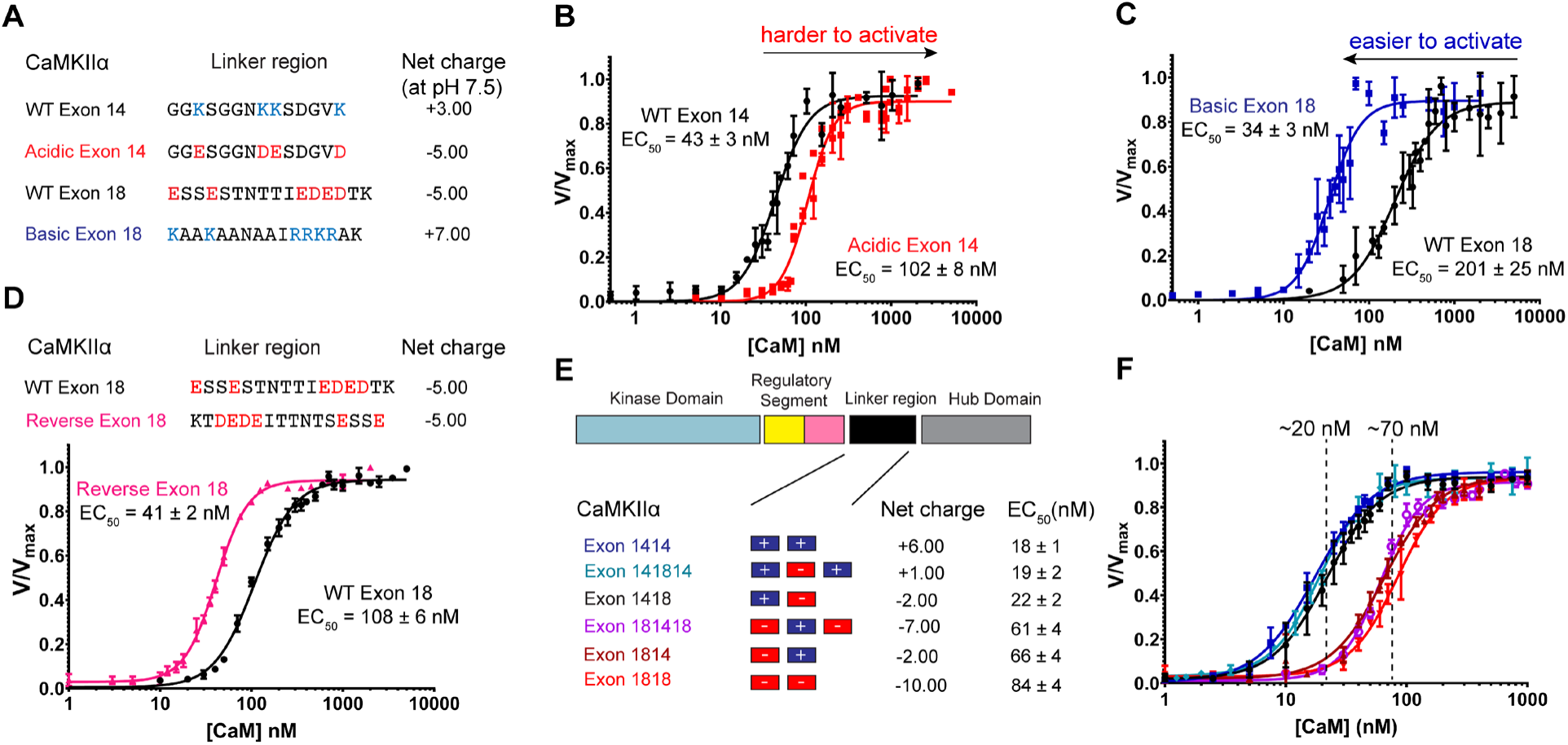
Positions of charged residues on the linker dictate Ca^2+^/CaM sensitivity. (A) Amino acid sequences of exon 14, exon 18, and the charge-reversal mutants. Basic and acidic residues that are mutated are colored blue and red, respectively. (B-C) CaMKII activity against syntide-2 is tested using linker mutants varying net charge. (D) CaMKII activity against syntide-2 is tested using WT exon 18 and the reverse sequence of exon 18. (E) Cartoon representing combinations of exon 14 and exon 18 used as the CaMKIIα variable linker region. Net charges and EC_50_ values from activity assays shown in (F) are listed. Each kinase assay comparison was done on the same day. Each data point of each graph represents the mean ± SD, n=3.

### The position of charge on the variable linker regulates Ca^2+^/CaM sensitivity

We next tested the effect of net charge on Ca^2+^/CaM binding. All net charges were calculated at pH 7.5. Exon 14 is highly basic (net charge +3) and exon 18 is highly acidic (net charge −5) (**Fig 2A**). If the net charge is driving this effect, then the inclusion of a basic exon makes it easier to activate CaMKII, while the inclusion of an acidic exon would make it harder to activate CaMKII. We acidified exon 14 by mutating all basic residues to acidic residues (**Fig 2A**). We hypothesized that including acidic exon 14 would make it harder to activate CaMKII. Indeed, acidic exon 14 has a ∼2-fold right-shifted EC_50_ (102 ± 8 nM) compared to WT exon 14 (43 ± 3 nM) (**Fig 2B**). Similarly, we basified exon 18 and this resulted in a ∼6-fold left-shifted EC_50_ (34 ± 3 nM) compared to the WT exon 18 (201 ± 25 nM) (**Fig 2C**). To test whether the charge position has an effect, we created a mutant where the sequence of exon 18 was reversed. This mutant left-shifted the EC_50_ value ∼2-fold (41 ± 2 nM) compared to WT exon 18 (108 ± 6 nM) (**Fig 2D**), indicating that the position of the charges does impact the accessibility of the regulatory segment to Ca^2+^/CaM. To further interrogate this position effect, we compared six combinations of exons 14 and 18 within CaMKIIα (**Fig 2E**). For reference, CaMKIIα with exons 14 and 18 included [CaMKIIα (14,18) or CaMKIIα-30] is the most prevalent CaMKII variant in the brain ^10^. CaMKIIα mutants with basic exon 14 positioned proximal to the regulatory segment have left-shifted EC_50_ values (∼20 nM), whereas all mutants with acidic exon 18 in this position have right-shifted EC_50_ values (∼70 nM) (**Fig 2F**).

### The linker sequence affects the compactness of the holoenzyme

We used small-angle x-ray scattering (SAXS) which has previously been implemented to study conformational differences in CaMKII variants ^6,10,24^. We performed SAXS coupled with size exclusion chromatography and multi-angle scattering and observed a single elution peak for each sample (**Figs. 3A, B**). Guinier analyses (**Figs. 3C, D**) revealed a significant difference in the radius of gyration (ANCOVA, F_1,38_ = 173.86, p <0.001) between α exon 18 (R_g_ = 60 ± 1 Å) and α exon 14 (R_g_ = 68 ± 2 Å). In addition, the maximal dimensions derived from pair-wise distribution functions also showed a substantial difference between α exon 18 (D_max_ = 190 Å) and α Exon 14 (D_max_ = 257 Å) (**Fig. 3E**). These data suggest that inclusion of α exon 18 results in more compact holoenzymes compared to α exon 14-containing holoenzymes.

**Figure 3.**
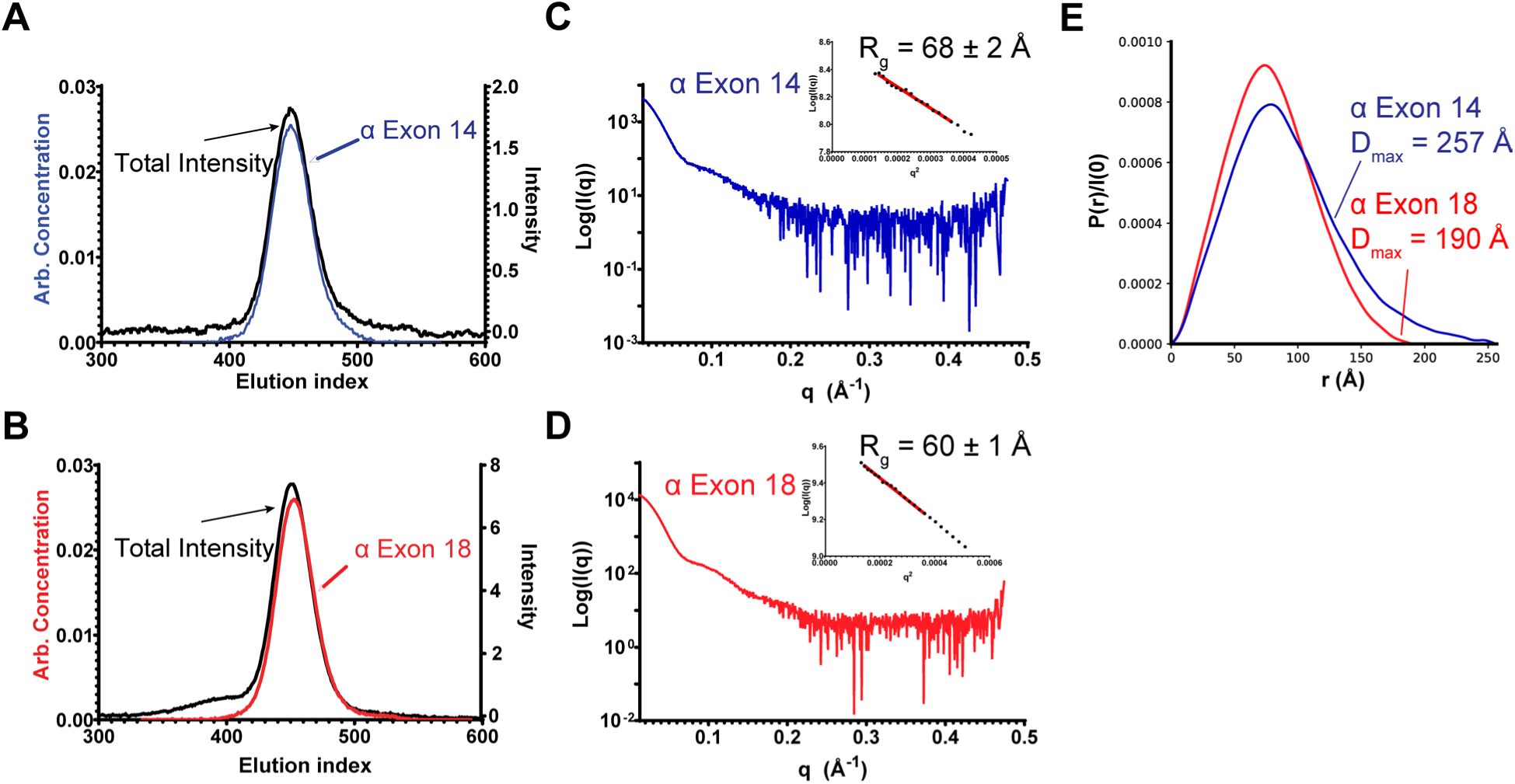
SEC-SAXS reveals a significant shape difference between CaMKIIα containing exon 14 or exon 18. (A-B) Size-exclusion chromatographs of CaMKIIα exon 14 and exon 18 are shown. The protein was monitored with absorbance at 280 nm as shown in black trace. Evolving factor analysis (EFA) indicates the main eluent component under the peak, which is shown for exon 14 (blue) and exon 18 (red). (C-D) SAXS profiles of CaMKIIα exon 14 and exon 18. Insets are the Guinier analysis to interpolate the R_g_ value ± SD of the fit for each construct. (E) Pair distance distribution of SAXS profiles of CaMKIIα exon 14 and CaMKIIα exon 18 were calculated in GNOM program in RAW v.2.1.4.

We next asked whether there were any structural differences between α exon 14 and α exon 18 in the presence of Ca^2+^/CaM. Each protein was incubated with an increasing concentration of Ca^2+^/CaM and analyzed by SAXS. We observed that both α exon 14 and α exon 18 holoenzymes became more open in the presence of increasing Ca^2+^/CaM concentrations (**Fig. S2**). The R_g_ values of α exon 14 were consistently higher than α exon 18 until they both reached ∼99 Å with saturating Ca^2+^/CaM, suggesting a similar extended conformation for both α exon 14 and α exon 18.

### Exon 14 and 18 containing CaMKIIs have different Ca^2+^ responsiveness in the cell

To examine the effect of the linker sequence on CaMKII holoenzyme conformation in a physiologically relevant setting, we employed a FRET-based assay using variants of the biosensor Camui ^8,25–27^. The Camui sensor is comprised of CaMKII with a FRET pair fused to the N- and C-termini. This sensor exploits the large conformational change that CaMKII undergoes when bound to Ca^2+^/CaM. Camui is also sensitive enough to report a FRET difference between Ca^2+^/CaM-bound state and phosphor-Thr286 state^28^. To compare CaMKIIα exon 14 to exon 18, we created two separate Camui variants containing either exon 14 or exon 18 as the linker region. CaMKIIα was fused to monomeric super enhanced CFP at the N-terminus and mCitrine YFP at the C-terminus (**Fig. 4A, 4B**). Changes in the YFP/CFP ratio report on the activated (open state, low FRET) and unactivated (compact state, high FRET) of the holoenzyme.

**Figure 4.**
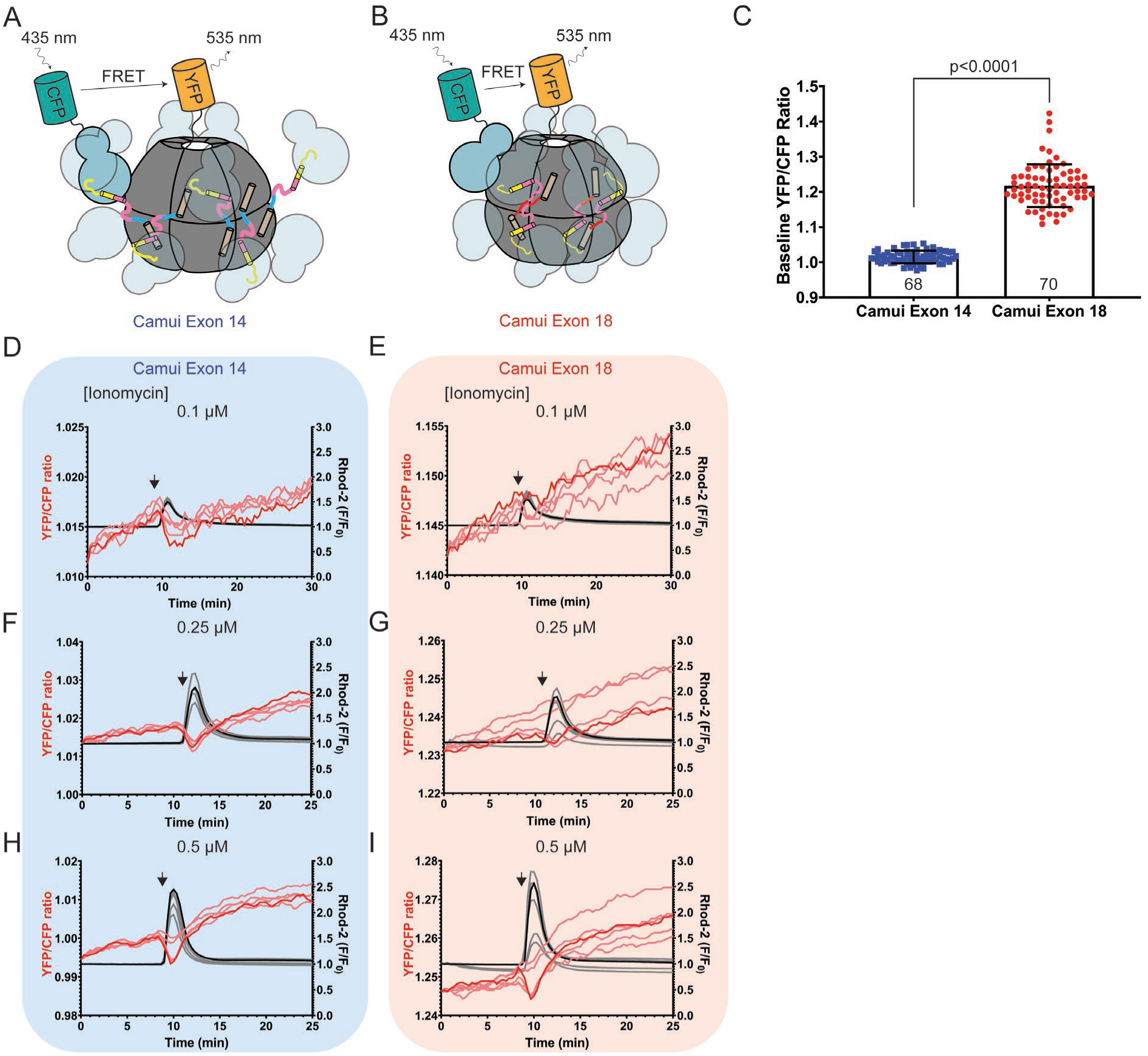
A FRET-based assay in mouse oocytes reports a change in Ca^2+^ responsiveness comparing CaMKIIα exon 18 to CaMKIIα exon 14. (A, B) A model of Camui exon 14 (A) and Camui exon 18 (B) that contains an N-terminal mseCFP and a C-terminal YFP (mCitrine). For clarity, only one pair of CFP and YFP is shown. (C) Baseline FRET signal within the first minute of oocytes expressing either Camui exon 14 or Camui exon 18. The number within each bar is the total number of oocytes per condition. Data is shown as mean ± SD. (D-I) YFP/CFP ratios (reporting on conformational changes) and Rhod-2 traces (reporting on cytosolic Ca^2+^ level) in mouse eggs expressing Camui exon 14 (D, F, H) and expressing Camui exon 18 (E, G, I).

We used mouse oocytes as a system to study CaMKII responsiveness as it has been done previously ^27^. We injected mRNAs encoding either Camui exon 14 or Camui exon 18 and simultaneously measured CaMKII activity (Camui FRET) and Ca^2+^ levels (rhodamine-2). We observed a statistically significant higher baseline YFP/CFP ratio in Camui exon 18 compared to Camui exon 14 (**Fig. 4C**, two-tailed t-test, t=23.08, df=45, p<0.0001). This result is consistent with our SAXS data showing CaMKIIα exon 18 is more compact than CaMKIIα exon 14.

We asked if the different conformations of the holoenzyme would affect how it responds to Ca^2+^ signaling. We used ionomycin to induce Ca^2+^ release in mouse oocytes. Previously, Camui with a 30-residue linker expressed in mouse oocytes reported different levels of activity upon varying levels of cytosolic Ca^2+^ by exposing oocytes to different concentrations of ionomycin ^27^. Adopting this strategy, we exposed oocytes expressing either Camui exon 14 or Camui exon 18 to different concentrations of ionomycin and monitored the emission changes. As expected, both Camui exon 14 and Camui exon 18 showed increasing levels of emission changes increases with higher ionomycin concentration, however the responsiveness between variants was different (**Fig 4D-I**). At 0.1 µM of ionomycin, we observed a clear change in YFP/CFP ratio for Camui exon 14, however the change for Camui exon 18 was barely detectable despite similar levels of Ca^2+^ release (**Fig. 4D, 4E**). We observed a similar result at 0.25 µM ionomycin where Camui exon 14 clearly responded to the rise of Ca^2+^ level while Camui exon 18 had a comparably smaller response (**Fig. 4F, 4G**). Finally, at 0.5 µM of ionomycin both Camui exon 14 and Camui exon 18 responded similarly under this saturated condition (**Fig. 4H, 4I**). Overall, our biochemical and cellular data support a model where exon 18 renders a more compact CaMKII holoenzyme that causes reduced Ca^2+^/CaM sensitivity.

### The CaMKII holoenzyme is comprised of domain-swapped dimers

We determined the X-ray crystal structure of no-linker full-length human CaMKIIδ-0 (residues 10-447) to 2.3 Å (**Table 1**). The crystallization construct contained one mutation (D336C), introduced for labeling experiments, which is surface exposed and not involved in crystal packing in the reported lattice. The structure was resolved to contain three CaMKII subunits in the asymmetric unit, which generates a dodecameric holoenzyme by the crystallographic P4_2_22 symmetry (**Fig. 5A, Fig. S3**). The overall architecture is identical to that of CaMKIIα with no linker (PDB: 3SOA ^6^). However, the improved resolution and the different space group allowed for unambiguous building of residues 300-313, which includes the junction between the regulatory segment and hub domain herein deemed the ‘connector segment’ (**Fig. S3A**). The electron density in this region extends away from the kinase domain of one subunit toward the hub domain of the opposite stacked ring, resulting in a domain-swapped dimer with the kinase domain of one subunit associates with the hub domain of a copy across a molecular 2-fold axis (**Fig. 5A, B**). The CaM footprints of the domain-swapped dimer run antiparallel to one another and are in extensive contact (**Fig. 5B**).

**Figure 5.**
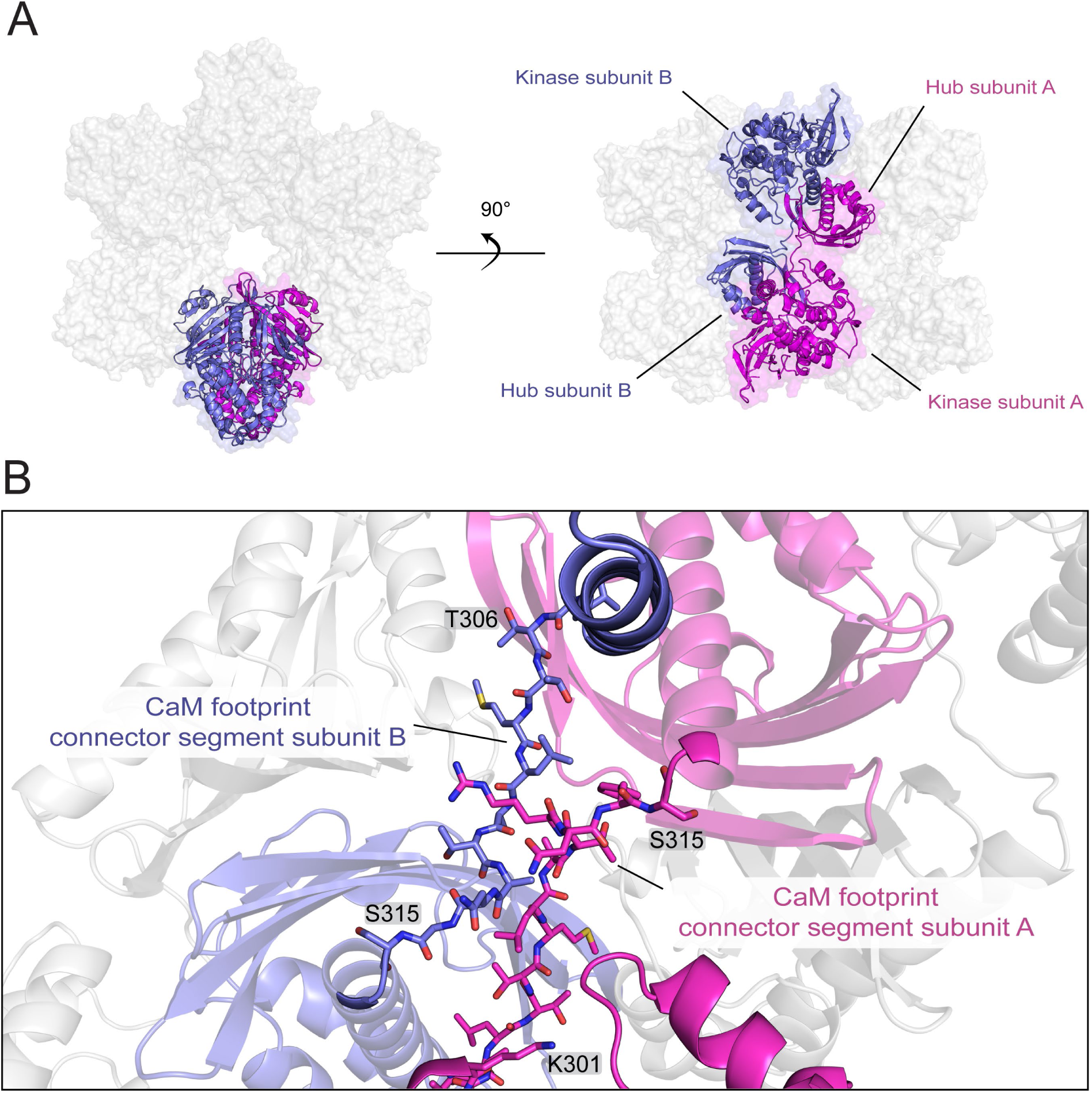
The CaMKIIδ-0 holoenzyme is comprised of domain-swapped dimers. (A) Surface representation of CaMKIIδ holoenzyme crystal structure (PDB code: 8USO) in two perspectives showing the interaction between the kinase domain of one subunit and the hub domain of another subunit. (B) The connector segments (defined as residues 305-314) in the CaM footprints of two domain-swapping subunits are shown as sticks. The kinase domain of subunit B has been omitted for clarity.

**Table 1.** Data Collection and Refinement Statistics.

| **Data collection** |  |
| --- | --- |
| CaMKIIδ holoenzyme (PDB: 8USO) | |
| Collection wavelength: 0.979 Å |  |
| Space group | $P4_22$ |
| Cell dimensions |  |
| a, b, c (Å) | 174.984, 174.984, 140.416 |
| $\alpha$, $\beta$, $\gamma$ (°) | 90, 90, 90 |
| Resolution range (Å) | 48.53 – 2.30 (2.34 – 2.30) |
| Unique reflections | 96,528 (4,754) |
| $R_{\text{merge}}$ | 0.158 (1.626) |
| $R_{\text{meas}}$ | 0.162 (1.691) |
| $R_{\text{pim}}^a$ | 0.036 (0.446) |
| Mean $I/\sigma(I)$ | 23.5 (1.4) |
| Completeness (%) | 100.0 (99.9) |
| Redundancy | 20.0 (12.9) |
| $CC_{1/2}$ | 0.992 (0.611) |
| $CC^*$ | 0.998 (0.871) |
| Wilson $B$-value (Å<sup>2</sup>) | 31.3 |
| **Refinement** |  |
| Resolution (Å) | 48.53 – 2.30 (2.36 – 2.30) |
| No. reflections $R_{\text{work}} / R_{\text{free}}$ | 86,683/2,000 (2,630/62) |
| Completeness (%) | 89.8 (40.0) |
| $R_{\text{work}}$ (%) | 18.4 (21.6) |
| $R_{\text{free}}$ (%) | 21.3 (28.7) |
| Atoms (non-H protein/ions/water) | 10,457/21/302 |
| Ramachandran plot (%) (favored/additional/disallowed) | 94.9/4.6/0.5 |
| Mean $B$-value (Å<sup>2</sup>) (chains A/B/C/ions/water) | 54.4/47.3/51.0/35.0/38.2 |
| R.m.s.d. bond length (Å) | 0.002 |
| R.m.s.d. bond angle (°) | 0.73 |
| Clashscore/Overall score<sup>b</sup> | 3.21/1.46 |
| Maximum likelihood coordinate error | 0.23 |

High-resolution bin statistics are shown in parentheses.
<sup>a</sup> $R_{\text{pim}} = 100 \sum_h \sum_i [1/(n_h - 1)]^{1/2} |I_{h,i} - \langle I_h \rangle| / \sum_h \sum_i \langle I_{h,i} \rangle$ , where $n_h$ is the number of observations of reflections $h$ .
<sup>b</sup> As defined by the validation suite MolProbity <sup>67</sup>.

The hub:hub interfaces that drive oligomerization are equatorial and lateral interfaces with extensive hydrophobic interactions and hydrogen bonding (**Fig. 6A**). We rationally designed interfacial mutations to validate the structure and to drive oligomer dissociation. We compared fresh preparations of full-length CaMKIIδ constructs with no variable linker region (our crystallization construct) to mutations at the lateral and/or equatorial hub interfaces and employed mass photometry to measure the masses of single particles.

**Figure 6.**
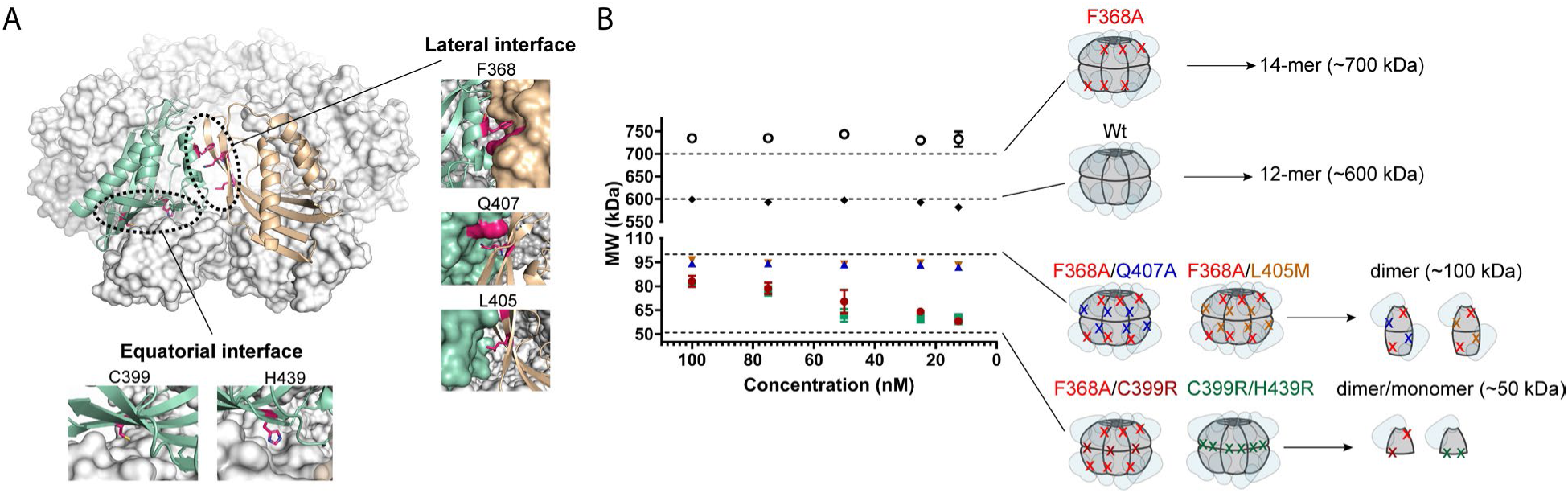
Two mutations are necessary to form predominant dimer or monomer populations. (A) The dodecameric CaMKIIδ hub from the crystal structure presented here is shown with the kinase domains omitted for clarity. Residues highlighted on the right (F368, L405, Q407) are positioned in the lateral hub interface and on the bottom (C399 and H439) are positioned in the equatorial hub interface. (B) Mass photometry analyses at different concentrations of CaMKIIδ holoenzymes with mutations at either or both hub interfaces. Cartoon representations on the right show the positions of the mutations (indicated as X). The kinase domains are translucent for clarity. Data are represented as mean molecular weight ± SD, n=3.

WT CaMKIIδ holoenzymes form dodecameric oligomers (600 kDa) from 100 nM to 12.5 nM (the lowest concentration we can reliably detect), which was consistent with our crystal structure (**Fig. 6B**, **Fig. S4**). In the WT sample, we tested F368A (F367A in CaMKIIα numbering), which has previously been shown to destabilize the oligomer ^7^. We found that F368A CaMKIIδ holoenzymes formed predominantly tetradecameric oligomers (700 kDa) (**Fig. 6B**). At 100 nM, we observed ∼10% of the total population as dimers (100 kDa) and ∼5% as tetramers (200 kDa) (**Fig. S4**). This percentage of dimers increased to ∼20% at 25 nM. Compared to WT, which had negligible smaller species across the concentration range tested (<7% monomer/dimer), F368A destabilized the oligomer to form dimers at lower concentrations.

To further promote a dimeric population, we introduced a second mutation to complement F368A (lateral mutation). Both L405 and Q407 also pack against the neighboring hub subunit along the lateral interface (**Fig. 6A**). The double mutants (F368A/L405M and F368A/Q407A) produced a dimeric population (100 kDa) from 100 nM to 12.5 nM (**Fig. 6B**, **Fig. S4**). There was ∼5% tetradecameric holoenzyme (700 kDa) at 100 nM, but this disappeared below 75 nM (**Fig. S4**). Finally, to promote a monomeric population, we introduced mutations at the equatorial interface (C399R and H439R). The double mutant C399R/H439R produced a population between dimer and monomer (∼80 kDa) at concentrations above 75 nM, and produced monomers below 50 nM (**Fig. 6B**, **Fig. S4**). There were also 300 kDa populations (6-mers) in the C399R/H439R sample above 75 nM, but these dissociated below 50 nM with a concomitant increase in the monomeric population (**Fig. S4**).

The domain-swapped conformation suggests that the interaction between the hub domain and the kinase domain could stabilize the dimer across the equatorial plane. To test this, we introduced C399R (equatorial mutation) to the F368A (lateral mutation) background to disrupt both interfaces while leaving the kinase-hub interface intact. Unlike the H439R/C399R mutant, the resulting F368A/C399R mutant did not contain any high molecular weight populations (**Fig. 6B**, **Fig. S4**). The molecular weight distribution of F368A/C399R skewed toward 80 kDa, suggesting an equilibrium between monomers and dimers at concentrations above 75 nM (**Fig. 6B, Fig. S4**). Similar to the H439R/C399R mutant, the F368A/C399R mutant showed predominantly monomers at concentrations lower than 50 nM (**Fig. 6B**, **Fig. S4**). These results suggest that the kinase-hub interactions contribute to holoenzyme formation, and it dissociates with hub dissociation.

We next determined the functional implications of oligomerization on CaMKII activation. We measured the activity of CaMKIIδ oligomer mutants at varying Ca^2+^/CaM concentrations towards syntide-2, as has been done previously ^6,10,29^. We observed that all CaMKIIδ constructs at final concentration of ∼13 nM exhibited positive cooperativity ranging from 1.7 to 2 regardless of the oligomeric state (**Fig. S5**). The EC_50_ values for the CaMKIIδ mutants were left-shifted compared to the WT (**Fig. S5**). The dimeric mutants (F368A/L405M and F368A/Q407A) and monomeric mutants (F368A/C399R and C399R/H439R) had a 2-fold lower EC_50_ value compared to WT CaMKIIδ (EC_50_ = 80 ± 4 nM). The tetradecameric mutant (F368A) showed an intermediate sensitivity toward Ca^2+^/CaM (EC_50_ = 61 ± 4 nM) relative to the other mutants and the wild type. These results demonstrate that the oligomeric state of CaMKII does tune Ca^2+^/CaM sensitivity.

### The domain-swapped dimer conformation orients electrostatic interactions by the linker region to influence autoinhibition

There are six basic residues in the CaM binding region of the regulatory segment (CaMKIIδ residue number, K^292^KFNA<u>R</u>^297^<u>RK</u>L<u>K</u>^301^), four of which (underlied) are involved in electrostatic interactions with Ca^2+^/CaM ^30^. We hypothesized that the negatively charged residues of the linker region could interact with the positively charged residues of the calmodulin binding region, which would reduce the affinity of calmodulin binding. To interrogate this, we employed molecular dynamic (MD) simulations paired with experimental validation by mutagenesis. We first modeled holoenzymes without a linker region or with linker regions comprised of WT exon 14, WT exon 18, or reversed exon 18. These were all modeled in the domain-swapped conformation and included two kinase domains and four hub domains to account for all interfaces (**Fig. S6**). The complexes remain intact throughout the 200-ns simulation (see Methods, **Fig. S7**). While the root-mean-squared fluctuation (RMSF) of all hub domains showed low magnitudes (**Fig. S8**), the RMSF of the kinase domain and regulatory segment from exon 14 and reversed exon 18 constructs are larger compared to those with no linker or WT exon 18 (**Fig. S8**). This indicates that there is more flexibility in the regulatory segment when WT exon 14 or reversed exon 18 is incorporated whereas there is less flexibility with the no linker or WT exon 18 constructs. This is consistent with a model that a more compact structure (WT exon 18) is harder to activate (see **Fig. 2**). We also ran simulations of these same linker constructs in the non-domain-swapped conformation (PDB: 3SOA), and there were no significant differences in RMSF throughout the structure between different linker constructs (**Fig. S9**).

In-depth pairwise residue interaction analyses indicated that the N- and C-terminal negative charges of exon 18 are in proximity to Lys 301 and Arg 297, which are involved in Ca^2+^/CaM binding ^30^. The negative charges on the N-terminus of exon 18 may contribute more because they were <5 Å away from Lys 301 and Arg 297 over the 200-ns trajectory (**Fig. 7A-D**, **Fig. S10**). To test the contribution of the N-terminal negative charges, we introduced EΔQ and DΔN mutations to exon 18 (**Fig. 7D**). Removing two negative charges in the N-terminus of exon 18 reduced the EC_50_ by half, whereas removing two negative charges at the C-terminal end did not change the EC_50_ significantly. Pairwise analyses showed that the negative charges on the C-terminus of exon 18 had less frequent contact with the positive charges of the regulatory segment (**Fig. S11**). Mutating all acidic residues on exon 18 made the variant (exon 18 mut3) significantly more sensitive to Ca^2+^/CaM (**Fig. 7E**).

**Figure 7.**
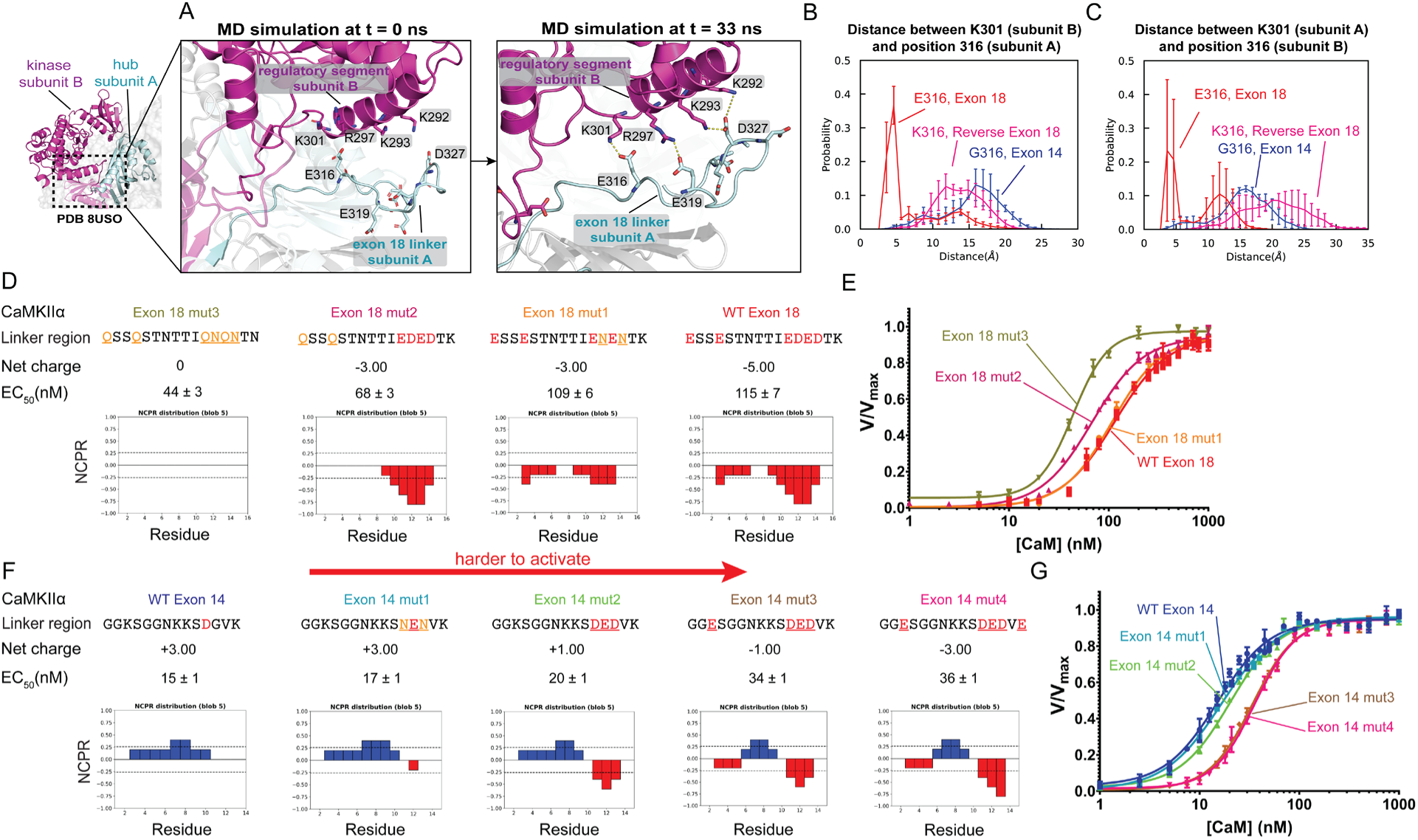
Specific electrostatic interactions between the linker and the calmodulin footprint drive the Ca^2+^/CaM sensitivity. (A) Snapshots from the MD simulation at time 0 and 33 ns containing the domain-swapped dimer (shown in inset) and exon 18 linker region. Electrostatic interactions are shown using dashed lines in the right panel. (B) Distributions of distance between K301 of subunit B and position 316 of subunit A from simulations of constructs with different linkers as labeled. (C) Same as B except comparing K301 of subunit A and position 316 of subunit B. See also Supplemental Figure 6-11. (D) Sequences of exon 18 mutants are shown to test the position of neutral or charged residues. Net charge and EC_50_ values (see panel E) are shown. (F) Sequences of exon 14 mutants are shown to test the position of neutral or charged residues. Net charge and EC_50_ values (see panel G) are shown. The net charge per residue (NCPR) graphs were plotted using CIDER server (http://157.245.85.131:8000/CIDER/). Data are shown as mean ± SD, n=3.

We next asked if a similar charge distribution can be applied to other linker sequences. Exon 14 has positive charges distributed across the sequence. We gradually increased the negative charge at the C-terminal portion of exon 14 (**Fig 7F**), which resulted in a negligible change in Ca^2+^/CaM sensitivity. Then, we mutated the first Lys of exon 14 to Glu (exon 14 mut3) and observed a significant increase in Ca^2+^/CaM sensitivity relative to WT exon 14 (**Fig. 7G**). Adding more negative charge to the C-terminus did not further right-shift the EC_50_ (exon 14 mut4) (**Fig. 7G**). These observations are consistent with the mechanism that the domain-swapped conformation of CaMKII positions the N-terminal negative charges of the linker to interact with positively charged residues of the calmodulin footprint, therefore regulating the accessibility of the regulatory segment. Negative charges at the C-terminal end had a more modest effect on sensitivity regulation.

## DISCUSSION

The structure of CaMKIIδ-0 provides the first evidence for a domain-swapped conformation within the holoenzyme. The previous structure of CaMKIIα-0 (PDB: 3SOA)^6^ had poor resolution in the connector segment. We had higher resolution of the residues that connect the kinase to the hub domain (CaMKIIα 299-304 and 309-314), which allowed us to unambiguously determine the domain-swapped conformation. Additionally, our structure provides three independent views of the connector segment from the three chains in the asymmetric unit (A, B, C, **Fig. S3A**). Each chain showed electron density that is only consistent with a domain-swapped dimer. The domain-swapping results in residues 307-315 running antiparallel to one another. Although we achieved higher resolution, the B-factors for residues 310-313 are relatively high (∼93 to 143 for Cα), likely indicating more flexibility in this region as might be expected for a junction between domains (**Fig. S3B**). Many domain-swapped oligomers contain flexible linkers to accommodate this conformation ^31,32^. Further studies will be necessary to determine whether domain swapping occurs in all CaMKII structures or whether there are dynamic fluctuations between swapped and unswapped conformations. The data presented here support a domain-swapped model.

A domain-swapped conformation of CaMKII has several implications. As tested here, the orientation of the linker region to the Ca^2+^/CaM binding site provides an opportunity for the linker to regulate the accessibility to Ca^2+^/CaM. Domain-swapping increases the number of communication points between subunits, which in turn, provides a larger landscape for allosteric effects such as cooperativity ^29^ and hub regulation of sensitivity ^6,10^. In comparing the compactness (**Fig. 3**) to conformational fluctuations (**Fig. 4C**), we find that the more compact exon 18 exhibits a higher variance in basal FRET signal, whereas the more extended exon 14 shows a lower variance. We interpret this to mean that the compact structure fluctuates between hub-bound and hub-unbound while the extended structure rarely adopts a hub-bound state. In addition, rationally designed mutations to disrupt hub interfaces yield CaMKII holoenzymes with different oligomeric states and Ca^2+^/CaM sensitivities (**Fig. 6, Fig. S5**). These mutations can be used to create a stable dimeric population (F368A/Q407A, F368A/L405M) or monomeric population under 50 nM (F368A/C399R, C399R/H439R). From these experiments, it is clear that stoichiometry affects CaMKII activation profiles, providing impetus to fully understand the relationship between CaMKII structure and Ca^2+^ sensitivity.

We provide a model using the domain-swapped structure in which specific linker residues interact with the regulatory segment, ultimately tuning the affinity of Ca^2+^/CaM. In 2000, Kwiatkowski and McGill showed that substituting a basic residue with an acidic one in the first linker exon of CaMKIIγ (R318E) increased the EC_50_ for Ca^2+^/CaM, consistent with our results ^33^. Our study aimed to make direct comparisons by using linker exons that are conserved across paralogs (combinations of exons 14 and 18). However, it is important to note that some of the CaMKIIα variants we tested, containing only exon 14 or exon 18, have not been observed in cells ^10,17,34^. CaMKIIβ with exon 18 only and CaMKIIδ with exon 14 only have been detected in adult human hippocampus^10^. Nevertheless, our comparisons allowed us to build a model where sensitivity is dependent on the distribution of charged residues in the linker, specifically the linker region most proximal to the regulatory segment (**Fig 2**). The model we provide fits the data for the set of variants we have tested here, however, because of the enormous diversity of CaMKII variants due to alternative splicing, we may need to expand this model to comprehensively explain the intricate dynamics of the autoinhibited state ^35^. Because we have concluded that the linker region most proximal to the regulatory segment is sufficient for influencing sensitivity, it may only be necessary to consider this part of the linker. Future efforts will focus on testing a broader range of variants to refine this model.

Charged residues in the CaMKII linker have been implicated in human disease. Researchers have identified K323N or K328E mutations in CaMKIIα linker in patients with dementia ^36^. These mutants were found to impair Ca^2+^/CaM binding and autophosphorylation at T286, resulting in reduced CaMKII binding to NR2B subunit of the ionotropic NMDA receptor, which is consistent with our findings here ^36^. Additionally, another study demonstrated that phosphorylation at S331 in CaMKIIα effectively inhibited CaMKII and led to an enhancement of memory-based addiction treatment outcomes ^37^. Indeed, CaMKIIα S331E had decreased activity compared to WT or S331A ^37^. While our mass spectrometry analysis detected phosphorylation at S330, phosphorylation on exon 18 was insufficient to explain our observed shift in Ca^2+^/CaM sensitivity (**Fig. S1G**). Previous research has also suggested that the charge of the linker region, whether through phosphorylation or insertion of charged residues by alternative splicing, influences localization and interaction with binding partners ^38–40^. Future studies will investigate the role of linker phosphorylation in CaMKII sensitivity, while also considering the potential impact of other post-translational modifications such as methylation and ubiquitination.

CaMKII structure and regulation are intimately linked. The CaMKII holoenzyme adopts an autoinhibited state in the absence of Ca^2+^, and the strength of autoinhibition relies on the sequence of the hub domain ^10^ as well as the length ^6,8,10,19,41^ and distribution of charge of the linker region, as shown here. Recent studies suggest that understanding CaMKII regulation requires consideration of its non-enzymatic scaffolding role ^42,43^ in addition to its kinase activity. While our focus here has been on CaMKII activity, it is worth noting that this is also tied to its scaffolding function, which necessitates phosphorylation at T286.

Enhanced CaMKII activation may improve kinetics for scaffolding capabilities. This new domain-swapped perspective on the CaMKII holoenzyme structure necessitates re-interpreting biochemical and structural data.

CaMKII is required for signaling in neurons, oocytes, and cardiomyocytes – all cells that use Ca^2+^ oscillations for communication. The responsiveness of CaMKII to Ca^2+^ is dictated by its structure, where stronger autoinhibition decreases sensitivity to Ca^2+^. Our results add to this picture by now incorporating the sequence of the variable linker region as an additional regulatory component. While each cell likely expresses a distinct set of splice variants of CaMKII paralogs, there is no data on protein levels at single-cell resolution. Knowing the abundance of all expressed CaMKIIs would provide testable hypotheses on the cellular response to Ca^2+^ and its variations across different cells and tissues.

## METHODS

### Plasmid construction

cDNAs encoding full-length CaMKII variants were subcloned into a pSMT3 vector containing N-terminal 6×His followed by a SUMO tag. The DNA sequences of exon 14 and exon 18 contain incomplete codons for all CaMKII genes tested in this study. To clone exon 18, we remove the last nucleotide from the last exon (exon 12) of the regulatory segment immediately preceding the linker region, which results in no changes in the protein sequences of the regulatory segment and exon 18. To clone exon 14, we added one or two nucleotides to the 3’-end of exon 14 to keep the sequence in frame, coding for the native residue at the linker/hub junction: Val (CaMKIIα), Ala (CaMKIIβ), Ala (CaMKIIδ). The DNA sequences of exon 14 and exon 18 are shown below. Added nucleotides are underlined.

α Exon 14: GAGGGAAGAGTGGGGGAAACAAGAAGAGCGATGGTGTGAA<u>GG</u>

α Exon 18: GGAATCCTCAGAGAGCACCAACACCACCATCGAGGATGAAGACACCAAAG

β Exon 14: CGGCCAAGAGCCTGCTGAATAAGAAGGCCGACGGCGTGAAG<u>G</u>

β Exon 18: GAGAGCAGTGACAGCGCCAACACCACAATTGAGGATGAGGACGCCAAGG

δ Exon 14: CCGCCAAGAGCCTGCTGAAGAAACCGGATGGCGTGAAG<u>G</u>

δ Exon 18: GAAAGCACCGAGAGTAGCAATACCACCATCGAAGACGAGGATGTGAAGG

### Protein expression and purification

Rosetta 2(DE3)pLysS competent cells (Millipore) were used to recombinantly express all CaMKII variants as previously described ^10^. CaMKII variants were cotransformed with lambda phosphatase, and overnight expression was done by 1 mM IPTG (GoldBio) in 18 °C shaker. Cell pellets were resuspended and lysed in Buffer A [25 mM Tris-HCl (pH 8.5), 150 mM KCl, 50 mM imidazole, 10% glycerol; Sigma] with protease inhibitor mixture [0.2 mM AEBSF, 5.0 μM leupeptin, pepstatin (1 μg/ml), aprotinin (1 μg/ml), trypsin inhibitor (0.1 mg/ml), 0.5 mM benzamidine] and DNase (1 μg/ml) (Sigma). Protein purification was performed using ÄKTA pure chromatography system at 4°C. Modifications in the ionic strength and buffer pH were applied if necessary. After pelleting the cell debris, clarified cell lysate was loaded onto one or two 5-ml HisTrap FF NiNTA Sepharose column (GE) and eluted with a combination of 50% buffer A and 50% buffer B [25 mM tris-HCl (pH 8.5), 150 mM KCl, 1 M imidazole, 10% glycerol] for a final concentration of 0.5 M imidazole. Residual imidazole was separated from the protein using a HiPrep 26/10 Desalting column, and His SUMO tags were cleaved with Ulp1 protease overnight at 4°C in buffer C [25 mM Tris-HCl (pH 8.5), 150 mM KCl, 2 mM TCEP (GoldBio), 50 mM imidazole, 10% glycerol]. Cleaved tags were separated by a subtractive Ni-NTA step. Next, an anion exchange step was performed with a 5-ml HiTrap Q-FF or Mono Q™ 10/100 GL and protein was eluted with a KCl gradient. Eluted proteins were concentrated and further purified in gel filtration buffer [25 mM tris-HCl (pH 8.0), 150 mM KCl, 1 mM TCEP, 10% glycerol] using a Superose 6 Increase 10/300 size exclusion column. Pure fractions were then concentrated, aliquoted, flash-frozen in liquid nitrogen, and stored at −80°C until further use. All columns were purchased from GE.

Calmodulin (*Gallus gallus*) was recombinantly expressed from a pET-15b vector (a gift from A. Nairn, Yale School of Medicine) in BL21(DE3) cells (Millipore) and purified as previously described ^10,44^. The concentration of CaM stock was measured in triplicate by circular dichroism using Jasco J-1500 spectrophotometer scanning a wavelength spectrum between 260 and 205 nm. The characteristic wavelength of 222 nm was used in the calculation of calmodulin concentration as follows:

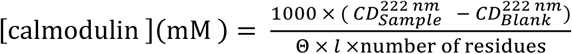

where the circular dichroism at 222 nm (CD^222nm^) is expressed in mdeg, Θ is the molar residual ellipticity (158000 deg cm^2^ dmol^-1^ residue^-1^), and *l* is the path length in cm.

### Couple Kinase Assay

Kinase activity was monitored by Synergy H1 microplate reader (BioTek) at 30 °C as previously described with minor modifications ^10,44^. Each reaction in the plate contained varying concentrations of calmodulin and a master mix solution with the following components at final concentrations: 25 mM Tris (pH 7.5; Thermo Fisher Scientific), 150 mM KCl (Sigma), Tris/MgCl_2_ buffer (50 mM/10 mM, respectively) (Thermo Fisher Scientific), 0.2 mM CaCl_2_ (Sigma), 1 mM phosphoenolpyruvate (Alfa Aesar), nicotinamide adenine dinucleotide (0.15 mg/ml; Sigma), pyruvate kinase (10 U/ml; Sigma), lactate dehydrogenase (30 U/ml; Millipore Sigma), 2 mM adenosine triphosphate (ATP) (Sigma), and 0.3 mM syntide-2 (LifeTein). The final pH of the reaction was 7.5-8, and the reaction was initiated with CaMKII for a final concentration of 13.3 nM (as monomer concentration). Absorbance of NADH was monitored at 15-s intervals for 10 min. The changes in the absorbance for each reaction over time were measured by calculating the slope of a five-point sliding window (1 min 15 s), and the maximum observed slope was used as the observed kinetic rate of that reaction. Kinetic rates across the calmodulin titration were fitted with Hill equation as shown below (Prism 8) to obtain the EC_50_ (defined as the calmodulin concentration needed to reach the half-maximal reaction velocity) and Hill coefficient values (h).

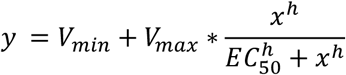

### Small-Angle X-ray Scattering

SAXS data were collected via mail-in service at the Advanced Light Source in Berkeley beamline 12.3.1. For SAXS analyses coupled with multi-angle light scattering in-line with size exclusion chromatography (SEC-SAXS-MALS), samples at concentrations from 3 to 10 mg/mL were prepared in SEC buffer [25 mM tris-HCl (pH 8.8), 250 mM KCl, 1 mM TCEP, 2% glycerol] and kept frozen in −20 °C until the run session. X-ray wavelength was set at λ=1.127 Å at a sample-to-detector distance of 2.1 m, resulting in q ranging from 0.011 to 0.473 Å^-1^ The scattering vector is q = 4πsinθ/λ, where 2θ is the scattering angle.

Size exclusion chromatography was performed using an Agilent 1260 series HPLC with a Shodex PROTEIN KW-804 column equilibrated in SEC buffer at a flow rate of 0.65 mL/min. Fifty five uL of each sample was applied to the SEC system and split 3-to-1 ratio between SAXS flow cells and a series of UV at 280 and 260 nm, multi-angle light scattering (MALS), quasi-elastic light scattering (QELS) and refractometer detectors. SAXS frames were collected continuously over the 30-min run at 3s exposures. Subsequent data analyses were performed in BioXTAS RAW ^45^ the ATSAS suit. The buffer regions were defined by the SAXS frames prior to the protein elution peak. The subtracted frames were analyzed by Evolving Factor Analysis ^46^ to distinguish co-eluted peaks prior to calculating R_g_ and D_max_. For SAXS experiment in the presence of Ca^2+^/CaM, 30 µM of CaMKII protein was incubated with 50 mM Tris (Thermo Fisher Scientific) pH 8.0, 0.2 mM CaCl_2_ (Sigma), 150 mM KCl (Sigma), 10 mM MgCl_2_ (Thermo Fisher Scientific), 2 mM ATP (Sigma), and varied CaM concentrations at room temperature for 1-2 minutes prior to X-ray exposure. The SAXS frame was collected after 10 min exposure. Data were collected using Rigaku’s BioSAXS-2000 SAXS instrument at UMass Core facility equipped with a P100K detector, MicroMax-007 HF X-ray source (λ=1.54 Å) and a sample-to-detector distance of 438.9 mm resulting in q ranging from 0.008 to 0.65 Å^-1^. The R_g_ was calculated for each of the subtracted frames using the Guinier approximation: I(q) = I(0) exp(−q^2^R_g_ ^2^/3) with the limits qR_g_ < 1.3.

### *In Vitro* Autophosphorylation of CaMKII

Purify full-length CaMKII (10 ug) was incubated for 10 minutes at room temperature (∼22 °C) in reaction buffer containing the following components at their final concentrations: 25 mM Tris (pH 8; Thermo Fisher Scientific), 150 mM KCl (Sigma), 1 mM TCEP (GoldBio), 5% glycerol (Thermo Fisher Scientific), 8 mM MgCl_2_ (Thermo Fisher Scientific), 2 mM ATP (Sigma), and 6 μM calmodulin. The reaction was quenched with 11 mM EDTA/ 11 mM EGTA then incubated with denaturation buffer (25 mM NH_4_HCO_3_, 8 M Urea, 10 mM DTT, pH 8) for 15 min at RT. Denatured sample was then alkylated by 20 mM iodoacetamide (1h, RT, dark) and trypsinized (2 ng/μL) overnight at 37 °C. Resulting peptides were mixed with 0.5% TFA, desalted using C18 Cartridge Solid Phase Extraction (Water Corporation), and dried using Speed Vac. Dried peptide was stored at −20 °C until further analysis.

### Mass Spectrometric Analysis

The dried peptide was resuspended in 0.1% TFA and resolved using Acclaim PepMap RSLC column C18 2 μm, 100 Å, 15 cm × 75 μm in an Easy nLC 1000 (Thermo Scientific). Solvent A consisted of 0.1% formic acid in water and solvent B, 0.1% formic acid in acetonitrile. Gradient elution profile (300 nL/min) was set as followed: 0 – 90 min, 0 – 35% B; 90 – 95 min, 35 – 95% B; 95 – 110 min, 95% B. Eluted peptides were analyzed on the Orbitrap Fusion mass spectrometer (Thermo Scientific) in data-dependent mode with dynamic exclusion enabled (60 s exclusion time). Full-scan spectra at resolution of 120K (at 200 m/z) were acquired in positive mode over the range of 200 – 1500 m/z, AGC target of 4e5, maximum injection time of 50 ms and 3 s cycle time. MS/MS scans were acquired in DDA mode by an orbitrap mass analyzer (HCD, collision energy 30%, resolution 30K, AGC target 5e5, maximum injection time 54 ms, isolation window 1.6 m/z). Thermo Raw files were analyzed in Proteome Discoverer 2.4.1 (Thermo Scientific).

### Mass Photometry

MP analyses were carried out with a OneMP mass photometer (Refeyn LTD, Oxford, UK) at room temperature as previously described with minor modifications ^47^. Glass coverslips (cat #630–2105) and gaskets (catalog #CW-50R-1.0) were cleaned with HPLC-grade water and isopropanol and dried under nitrogen gas before use. Gaskets are sealed onto coverslips during the drying process. Fifteen microliters of MP buffer (25 mM Tris pH 8.0, 150 mM KCl) was used to find the camera focus prior to loading the sample onto the gasket. Acquisition camera image size was set to “medium”. A 60-s movie was recorded for each sample in a 512 x 138 pixel at 1kHz frame rate with 0.95-ms exposure time. A standard curve was generated using thyroglobulin (MW 660 kDa), apoferritin (MW 440 kDa), BSA (66.5 kDa) with an error <1%. Proteins were purified and used on the same day to avoid freezing and thawing. The protein concentration was calculated from absorbance at 280 nm with extinction coefficients of 60850 M^-1^ cm^-1^ (CaMKIIδ) and 66350 M^-1^ cm^-1^ (CaMKIIα). Proteins were diluted from stock concentrations to 1 uM and stored on ice throughout the experiment. Immediately prior to a measurement, proteins were diluted to 400 nM, 300 nM, 200 nM, 100 nM, 50 nM and 5 uL at each concentration was applied to 15 uL of the MP buffer preloaded on the gasket to achieve final concentrations of 100 nM, 75 nM, 50 nM, 25 nM, 12.5 nM. Three replicates were performed for each protein at each concentration. Data collection was performed in AcquireMP v.2.3, and data were analyzed using DiscoverMP v2.3 software (Refeyn LTD, Oxford, UK).

### Crystallization and X-Ray Data Collection

Initial crystallization screening for the full-length CaMKIIδ (D336C) was done using the sitting vapor diffusion method at 20 °C with commercially available screening kits. The D336C mutation was made for another experiment involving chemical labeling the protein. The protein concentration was the same as the protein concentration (19 mg/ml) used to obtain CaMKIIα crystals. Three hits were further optimized by hanging drop vapor diffusion method. Final conditions are listed below. 20% glycerol was included in the cryo-solutions. Diffraction data were obtained at the Advanced Photon Source using Beamline 24-ID-E at 100 K with Eiger 16M detector. Each condition led to different crystal morphology, but they yielded the same crystal space group and unit cell values. Crystallization conditions:

1. 0.1 M pH 6.5 Bis-Tris, 16% PEG 2K, 0.2 M sodium malonate dibasic *this crystal was used for final structure determination and refinement.
2. 0.1 M MES pH 6.5, 1.6 M ammonium sulfate, 10% dioxane
3. 1.675 M Potassium phosphate dibasic/ Sodium phosphate monobasic pH 5.4, 3.35% isopropanol

### Data Processing and Structure Determination

All diffraction data were collected at beamline 19-ID (SBC-CAT) at the Advanced Photon Source (Argonne National Laboratory, Argonne, Illinois, USA) and processed in the program HKL-3000 ^48^ with applied corrections for effects resulting from absorption in a crystal and for radiation damage ^49,50^, the calculation of an optimal error model, and corrections to compensate the phasing signal for a radiation-induced increase of non-isomorphism within the crystal ^51,52^. The data displayed mild anisotropy. The structure was phased by molecular replacement in the program Phaser ^53^ using the coordinates of individual CaMKIIδ kinase (PDB: 2VN9) and hub domains (PDB: 2W2C) as search models. The unit cell volume was big enough to fit three copies of each domain. The regulatory segment and the region until hub domain manually built into clear electron density using Coot ^54^. Positional and isotropic atomic displacement parameter (ADP) as well as TLS ADP refinement was performed to a resolution of 2.30 Å for the model, using the program Phenix ^55^ with a random 2.3% of all data set aside for an R_free_ calculation. Model refinement in Phenix included automatic optimization of the X-ray terms to the stereochemistry and ADP weights, which resulted in an ideal balance between model stereochemistry (i.e., Ramachandran plot statistics, rotamer outliers, clashscores, etc.) and R_free_ values. The Ramachandran outliers in the model occur in regions of poor density, such as surface loops between secondary structural elements and the link between the kinase and hub domains. Data collection and structure refinement statistics are summarized in Table 1.

### Molecular Modeling and Simulation

#### Loop Modeling

To investigate linker dynamics and minimize computational cost, we chose to include two complete copies of monomers and only the two hub domains of the neighborly packed monomers from the dodecameric CaMKIIδ, which represents the minimal surrounding environment of interested linkers (**Fig. S6**). This construct preserves interactions from four hub domains surrounding two linkers. Overall, four models with different linkers (no linker, exon14, exon18 and reversed exon 18) and the missing loop residues (375-378) were generated using the loop modeling module in the MODELLER software ^56^. The built-in DOPE (Discrete Optimized Protein Energy) score in MODELLER was used to evaluate the energies of the final built models and then those optimized models were visually checked in VMD to assure all loops were closed in both ends with no steric clashes in the side chains.

#### Simulation protocol

The above four models were then individually solvated with TIP3P water using the CHARMM-GUI web server ^57^. The systems were neutralized and 200 mM KCl was added. The CHARMM36m force field was used, which has been optimized for both structured and intrinsically disordered proteins ^58^. The final four cubic boxes consist of ∼20,000 TIP3P water molecules, ∼470 ions, with a total atom of ∼210,000 and a cubic box of ∼13 nm in size. The energy minimization, equilibration and production were performed with GPU-accelerated GROMACS 2019.4 ^59^. Each system was first energy-minimized for 500 steps with the steepest descent method and then underwent a 125-ps equilibration simulation under NVT conditions at 300 K, with the protein backbone and sidechain atoms positionally restrained to the reference structure using harmonic force constants of 400 kJᐧmol^-1^ᐧnm^-2^ and 40 kJᐧ mol^-1^ᐧnm^-2^, respectively. In the production runs, we used a cutoff of 1.2 nm and a smooth switching function starting at 1.0 nm for calculation of non-boned interactions. The particle mesh Ewald (PME) method was used for long-range electrostatic interactions ^60^. The LINCS algorithm was used to constraint all hydrogen-involving bond lengths ^61^. The Nose-Hoover thermostat was used to maintain system temperature at 400 K ^62,63^. The production simulation of each system was carried out at 400 K under NPT conditions for 200 ns with two parallel runs, during which the Nose-Hoover barostat was used to maintain system pressure isotopically at 1 bar^64^. The temperature of 400 K was used to accelerate the simulation and better probe the difference among various constructs. No restraints were imposed during the production simulations at both 300 and 400K.

#### Analysis

To investigate the dynamics of linkers and kinase domains in each system, all trajectories were first aligned to their corresponding starting equilibrated structure using the C-alpha of all four hub domains. We then calculated RMSF profiles of all C-alpha atoms of the kinase domains, the regulatory segments, the linker regions and the hub domains. The RMSF was calculated by averaging the four copies of kinases in the 2 replicas and the error bars show the standard errors. To analyze interactions between the calmodulin-binding segment and the linkers, pairwise distances between C-alpha atoms of residues from the calmodulin-binding sequence (297R and 301K) and spatially adjacent residues from the linkers (the first and last residues of the linker sequence) were calculated. To find out the influence of the dynamics on the interaction between the kinase domain and hub domains, native contact analysis was performed. A cutoff of 4.5 Å was used to define whether a contact between heavy atoms of a pair of residues is formed. The total native contacts number is the number of contacts from the original structure, which was then normalized to be 100%. The native contacts number of each frame was calculated and normalized by the total native contacts number to get the native contact fraction.

### Animals and Collection of Mouse Oocytes

All animal procedures were performed according to research animal protocols approved by the University of Massachusetts Institutional Animal Care and Use Committee (IACUC, protocol #4579). All mice were housed in individually ventilated cages at 12-hour light/dark cycle with *ad libitum* access to food and water. C57BL/6J females were mated with DBA/2J male mice to generate BDF1 hybrids. 6- to 10-weeks-old female BDF1 mice were superovulated by intraperitoneal injections of 5 IU of pregnant mare serum gonadotropin (BioVendor) and 5 IU of human chorionic gonadotropin (hCG, Sigma) 48 hours after.

Metaphase-II stage oocytes were collected from the oviducts of these mice 13.5 h post-hCG injection in TL-HEPES media. Cumulus cells were removed with 0.26% bovine testes hyaluronidase (Sigma).

### mRNA Preparation and Microinjections

CaMKIIα exon 14 and CaMKIIα exon 18 were cloned into CKAR backbone ^65^ containing mseCFP and mCitrine by restriction cloning to make plasmids containing mseCFP-CaMKIIα Exon 14-mCitrine (Camui exon 14) and mseCFP-CaMKIIα Exon 18-mCitrine (Camui exon 18). Plasmids were linearized with DraIII-HF restriction enzyme overnight and mRNAs were in vitro transcribed using the mMESSAGE mMACHINE T7 Transcription Kit (Invitrogen), poly-adenylated using Poly(A) Tailing Kit (Invitrogen), and purified using MEGAclear Transcription Clean-up Kit (Invitrogen) according to manufacturer’s instructions. Purified mRNAs were diluted to 1 μg/μl concentration in nuclease-free water and stored at −80°C until use.

Microinjection of mouse oocytes was performed as described previously ^66^. Briefly, mRNAs were heated up to 95°C for 3 min, centrifuged at 12 100 g for 10 min at 4°C, and were loaded into glass micropipettes by aspiration. The plasma membranes of oocytes were penetrated using a piezo micro manipulator (PMAS-CT150, Prime Tech, Japan) and the mRNAs were delivered into the cytoplasm of oocytes by hydraulic pressure. Injected oocytes were placed in Advanced KSOM media (MR-101, Millipore Sigma) overlaid with mineral oil and allowed for translation up to 5 h at 37 °C in a humidified atmosphere of 5% CO2 until the time of monitoring.

### Ca^2+^ and FRET imaging

Imaging was performed as described before ^66^. Briefly, oocytes were loaded with 2.5 μM Rhod-2AM (Invitrogen) in TL-HEPES media supplemented with 0.02% Pluronic acid for 30 mins at RT. The oocytes were washed in protein- and nominally Ca^2+^-free TL-HEPES media (imaging media) and mounted on a glass-bottom imaging dish (MatTek Corp, Ashland, MA). The images were acquired every 20 seconds on three channels. YFP/CFP was performed using a CFP excitation filter, dichroic beam splitter, CFP and YFP emission filters (Chroma technology, Rockingham, VT; ET436/20X, 89007bs, ET480/40m, and ET535/30m), and Rhod-2 fluorescence was excited at 550 nm and recorded with a LP510 emission filter on an inverted Nikon Eclipse TE 300 microscope (Ghent, Belgium) equipped with a 20x objective, Photometrics SenSys CCD camera (Roper Scientific, Tucson, AZ), MAC5000 filter wheel/shutter control box (Ludl), and NIS-elements software (Nikon). Oocytes were treated with ionomycin (Tocris) at time points and concentrations specified in the figures. The averages of YFP/CFP ratios within the first minute of monitoring were used to compare the baseline YFP/CFP ratios. The average YFP/CFP ratios at time 0 in all oocytes expressing the same Camui per ionomycin concentration was calculated and set as the value for time 0 for all oocytes and applied to make the graphs. Rhod-2 values were calculated as actual value at X time/the average Rhod-2 value of the first minute of monitoring.

### Statistical Analysis

Statistical analysis of the data was accomplished using the GraphPad Prism 8 software. P-value < 0.05 was used as a predetermined threshold for statistical significance.

## DATA AVAILABILITY

The source data that support the findings of this study are available in Figshare with the identifier https://doi.org/10.6084/m9.figshare.27170322. The X-ray diffraction data has been deposited to the RCSB PDB (Accession code: 8USO). The shortgun proteomics data have been deposited to the ProteomeXchange Consortium via the MassIVE server with the dataset identifier MSV000094576 (doi:10.25345/C5057D414). SAXS data and models have been submitted to SASBDB (https://www.sasbdb.org/) with the following identifiers: CaMKIIα exon 14 (https://www.sasbdb.org/data/SASDVU7) SASDVU7; CaMKIIα exon 18 (https://www.sasbdb.org/data/SASDVV7) SASDVV7.

## CODE AVAILABILITY

CaMKII models and all scripts supporting the MD simulations can be found on Github (https://github.com/mdlab-um/camk2_linkers).

## ACKNOWLEDGEMENTS

We thank Dr. Luke Chao and Dr. Eric Strieter for their helpful comments on the manuscript. We thank Dr. Vincent Tagliabracci for assistance in obtaining beamtime. We thank Dr. Steve Eyles and the Mass Spectrometry core for helping with mass spectrometry samples. SAXS work was conducted at the Advanced Light Source (ALS), a national user facility operated by Lawrence Berkeley National Laboratory on behalf of the Department of Energy, Office of Basic Energy Sciences, through the Integrated Diffraction Analysis Technologies (IDAT) program, supported by DOE Office of Biological and Environmental Research. Additional support comes from the National Institute of Health project ALS-ENABLE (P30 GM124169) and a High-End Instrumentation Grant S10OD018483.

Crystallographic data shown in this report are derived from work performed at the Argonne National Laboratory, Structural Biology Center at the Advanced Photon Source. SBC-CAT is operated by UChicago Argonne, LLC, for the US Department of Energy, Office of Biological and Environmental Research under contract DE-AC02-06CH11357. This work was supported in part by NIH grant to M.M.S. (NIH R35GM145376), NIH grant to J.C. (NIH R35GM144045), a fellowship from the University of Massachusetts as part of the Chemistry-Biology Interface Training Program (National Research Service Award T32GM139789) to B.V.N., J.H., N.D., and A.P.T.-O, a fellowship from the University of Massachusetts as part of the NIH Biotechnology Traineeship (T32GM135096-01) to O.C.K.

## AUTHOR CONTRIBUTIONS

**Bao V. Nguyen:** Conceptualization; Investigation; Formal Analysis; Methodology; Validation; Visualization; Writing – original draft; Writing – review & editing. **Can Özden:** Conceptualization; Investigation; Formal Analysis; Methodology; Validation; Visualization; Writing – review & editing. **Kairong Dong:** Investigation; Formal Analysis; Methodology; Validation; Visualization; Writing – review & editing. **Oguz Koc**: Investigation; Formal Analysis; Methodology; Validation. **Ana P. Torres-Ocampo:** Resources; Writing – review & editing. **Noelle Dziedzic:** Conceptualization; Resources; Formal Analysis; Investigation. **Daniel Flaherty:** Resources; Investigation. **Jian Huang:** Methodology; Investigation; Resources. **Saketh Sankara:** Resources; Investigation. **Nikki Lyn Abromson:** Resources; Investigation. **Diana R. Tomchick**: Methodology; Resources; Formal Analysis. **Rafael Fissore**: Conceptualization; Methodology; Supervision. Writing – review & editing. **Jianhan Chen**: Conceptualization; Methodology; Funding acquisition; Supervision; Formal Analysis; Writing – review & editing. **Scott C Garman:** Conceptualization; Methodology; Supervision; Formal Analysis; Writing – review & editing. **Margaret M Stratton:** Conceptualization; Supervision; Formal Analysis; Visualization; Funding acquisition; Writing – original draft; Writing – review & editing.

## CONFLICT OF INTERESTS

The authors declare no conflict of interest.

## SUPPLEMENTAL INFORMATION

Document S1. Figures S1–S11

## Notes

### Competing Interest Statement

The authors have declared no competing interest.

### Summary of Updates

We have added cellular FRET experiments as well as new supplemental figures.

## REFERENCES

1. Backs, J. et al. The γ isoform of CaM kinase II controls mouse egg activation by regulating cell cycle resumption. Proceedings of the National Academy of Sciences 107, 81–86 (2010).

2. Gaido, O. E. R. et al. CaMKII as a Therapeutic Target in Cardiovascular Disease. Annual Review of Pharmacology and Toxicology 63, 249–272 (2023).

3. Nicoll, R. A. & Schulman, H. Synaptic memory and CaMKII. Physiological Reviews 103, 2897–2945 (2023).

4. Yang, E. & Schulman, H. Structural examination of autoregulation of multifunctional calcium/calmodulin-dependent protein kinase II. J Biol Chem 274, 26199–26208 (1999).

5. Hoelz, A., Nairn, A. C. & Kuriyan, J. Crystal Structure of a Tetradecameric Assembly of the Association Domain of Ca2+/Calmodulin-Dependent Kinase II. Molecular Cell 11, 1241–1251 (2003).

6. Chao, L. H. et al. A Mechanism for Tunable Autoinhibition in the Structure of a Human Ca2+/Calmodulin-Dependent Kinase II Holoenzyme. Cell 146, 732–745 (2011).

7. Bhattacharyya, M. et al. Molecular mechanism of activation-triggered subunit exchange in Ca2+/calmodulin-dependent protein kinase II. eLife 5, e13405 (2016).

8. Myers, J. B. et al. The CaMKII holoenzyme structure in activation-competent conformations. Nature Communications 8, 15742 (2017).

9. McSpadden, E. D. et al. Variation in assembly stoichiometry in non-metazoan homologs of the hub domain of Ca2+/calmodulin-dependent protein kinase II. Protein Science 28, 1071–1082 (2019).

10. Sloutsky, R. et al. Heterogeneity in human hippocampal CaMKII transcripts reveals allosteric hub-dependent regulation. Sci. Signal. 13, (2020).

11. Buonarati, O. R., Miller, A. P., Coultrap, S. J., Bayer, K. U. & Reichow, S. L. Conserved and divergent features of neuronal CaMKII holoenzyme structure, function, and high-order assembly. Cell Reports 37, 110168 (2021).

12. Küry, S. et al. De Novo Mutations in Protein Kinase Genes CAMK2A and CAMK2B Cause Intellectual Disability. The American Journal of Human Genetics 101, 768–788 (2017).

13. Stephenson, J. R. et al. A Novel Human CAMK2A Mutation Disrupts Dendritic Morphology and Synaptic Transmission, and Causes ASD-Related Behaviors. J Neurosci 37, 2216–2233 (2017).

14. Fujii, H. et al. Förster resonance energy transfer-based kinase mutation phenotyping reveals an aberrant facilitation of Ca2+/calmodulin-dependent CaMKIIα activity in de novo mutations related to intellectual disability. Front. Mol. Neurosci. 15, (2022).

15. Rigter, P. M. F. et al. Role of CAMK2D in neurodevelopment and associated conditions. The American Journal of Human Genetics 111, 364–382 (2024).

16. Chia, P. H. et al. A homozygous loss-of-function CAMK2A mutation causes growth delay, frequent seizures and severe intellectual disability. eLife 7, e32451 (2018).

17. Tombes, R. M., Faison, M. O. & Turbeville, J. M. Organization and evolution of multifunctional Ca2+/CaM-dependent protein kinase genes. Gene 322, 17–31 (2003).

18. Sloutsky, R. & Stratton, M. M. Functional implications of CaMKII alternative splicing. European Journal of Neuroscience 54, 6780–6794 (2020).

19. Bayer, K. U., Koninck, P. D. & Schulman, H. Alternative splicing modulates the frequency-dependent response of CaMKII to Ca2+ oscillations. The EMBO Journal 21, 3590–3597 (2002).

20. Duran, J., Nickel, L., Estrada, M., Backs, J. & Van Den Hoogenhof, M. M. G. CaMKIIδ Splice Variants in the Healthy and Diseased Heart. Front. Cell Dev. Biol. 9, 644630 (2021).

21. Franz, A. et al. Branch point strength controls species-specific CAMK2B alternative splicing and regulates LTP. Life Science Alliance 6, (2023).

22. Stratton, M. M., Chao, L. H., Schulman, H. & Kuriyan, J. Structural studies on the regulation of Ca2+/calmodulin dependent protein kinase II. Current Opinion in Structural Biology 23, 292–301 (2013).

23. Bhattacharyya, M., Karandur, D. & Kuriyan, J. Structural Insights into the Regulation of Ca ^2+^/Calmodulin-Dependent Protein Kinase II (CaMKII). Cold Spring Harb Perspect Biol 12, a035147 (2020).

24. Rosenberg, O. S., Deindl, S., Sung, R.-J., Nairn, A. C. & Kuriyan, J. Structure of the Autoinhibited Kinase Domain of CaMKII and SAXS Analysis of the Holoenzyme. Cell 123, 849–860 (2005).

25. Takao, K. et al. Visualization of Synaptic Ca ^2+^ /Calmodulin-Dependent Protein Kinase II Activity in Living Neurons. J. Neurosci. 25, 3107–3112 (2005).

26. Erickson, J. R., Patel, R., Ferguson, A., Bossuyt, J. & Bers, D. M. FRET-based sensor Camui provides new insight into mechanisms of CaMKII activation in intact cardiomyocytes. Circ Res 109, 729–738 (2011).

27. Ardestani, G., West, M. C., Maresca, T. J., Fissore, R. A. & Stratton, M. M. FRET-based sensor for CaMKII activity (FRESCA): A useful tool for assessing CaMKII activity in response to Ca2+ oscillations in live cells. J Biol Chem 294, 11876–11891 (2019).

28. Otmakhov, N., Regmi, S. & Lisman, J. E. Fast Decay of CaMKII FRET Sensor Signal in Spines after LTP Induction Is Not Due to Its Dephosphorylation. PLOS ONE 10, e0130457 (2015).

29. Chao, L. H. et al. Inter-subunit capture of regulatory segments is a component of cooperative CaMKII activation. Nat Struct Mol Biol 17, 264–272 (2010).

30. Rellos, P. et al. Structure of the CaMKIIδ/Calmodulin Complex Reveals the Molecular Mechanism of CaMKII Kinase Activation. PLOS Biology 8, e1000426 (2010).

31. Gronenborn, A. M. Protein acrobatics in pairs — dimerization via domain swapping. Current Opinion in Structural Biology 19, 39–49 (2009).

32. Rousseau, F., Schymkowitz, J. & Itzhaki, L. S. Implications of 3D Domain Swapping for Protein Folding, Misfolding and Function. in Madame Curie Bioscience Database [Internet] (Landes Bioscience, 2013).

33. Kwiatkowski, A. P. & McGill, J. M. Alternative Splice Variant of γ-Calmodulin-Dependent Protein Kinase II Alters Activation by Calmodulin. Archives of Biochemistry and Biophysics 378, 377–383 (2000).

34. Cook, S. G. et al. Analysis of the CaMKIIα and β splice-variant distribution among brain regions reveals isoform-specific differences in holoenzyme formation. Sci Rep 8, 5448 (2018).

35. Gaertner, T. R. et al. Comparative Analyses of the Three-dimensional Structures and Enzymatic Properties of α, β, γ, and δ Isoforms of Ca2+-Calmodulin-dependent Protein Kinase II *. Journal of Biological Chemistry 279, 12484–12494 (2004).

36. Wang, P. et al. PTENα Modulates CaMKII Signaling and Controls Contextual Fear Memory and Spatial Learning. Cell Reports 19, 2627–2641 (2017).

37. Rich, M. T. et al. Phosphoproteomic Analysis Reveals a Novel Mechanism of CaMKII Regulation Inversely Induced by Cocaine Memory Extinction versus Reconsolidation. Journal of Neuroscience 36, 7613–7627 (2016).

38. Heist, E. K., Srinivasan, M. & Schulman, H. Phosphorylation at the Nuclear Localization Signal of Ca2+/Calmodulin-dependent Protein Kinase II Blocks Its Nuclear Targeting*. Journal of Biological Chemistry 273, 19763–19771 (1998).

39. O’Leary, H., Lasda, E. & Bayer, K. U. CaMKII␤ Association with the Actin Cytoskeleton Is Regulated by Alternative Splicing□D. Molecular Biology of the Cell 17, 10 (2006).

40. Kim, K. et al. A Temporary Gating of Actin Remodeling during Synaptic Plasticity Consists of the Interplay between the Kinase and Structural Functions of CaMKII. Neuron 87, 813–826 (2015).

41. De Koninck, P. & Schulman, H. Sensitivity of CaM Kinase II to the Frequency of Ca ^2+^ Oscillations. Science 279, 227–230 (1998).

42. Tullis, J. E. et al. LTP induction by structural rather than enzymatic functions of CaMKII. Nature 621, 146–153 (2023).

43. Chen, X. et al. CaMKII autophosphorylation is the only enzymatic event required for synaptic memory. Proceedings of the National Academy of Sciences 121, e2402783121 (2024).

44. Stratton, M. et al. Activation-triggered subunit exchange between CaMKII holoenzymes facilitates the spread of kinase activity. eLife 3, (2014).

45. Hopkins, J. B., Gillilan, R. E. & Skou, S. *BioXTAS RAW*: improvements to a free open-source program for small-angle X-ray scattering data reduction and analysis. J Appl Crystallogr 50, 1545–1553 (2017).

46. Meisburger, S. P. et al. Domain Movements upon Activation of Phenylalanine Hydroxylase Characterized by Crystallography and Chromatography-Coupled Small-Angle X-ray Scattering. J. Am. Chem. Soc. 138, 6506–6516 (2016).

47. Torres-Ocampo, A. P., et al. Characterization of CaMKIIα holoenzyme stability. Protein Science 29, 1524–1534 (2020).

48. Minor, W., Cymborowski, M., Otwinowski, Z. & Chruszcz, M. HKL −3000: the integration of data reduction and structure solution – from diffraction images to an initial model in minutes. Acta Crystallogr D Biol Crystallogr 62, 859–866 (2006).

49. Borek, D., Minor, W. & Otwinowski, Z. Measurement errors and their consequences in protein crystallography. Acta Crystallogr D Biol Crystallogr 59, 2031–2038 (2003).

50. Otwinowski, Z., Borek, D., Majewski, W. & Minor, W. Multiparametric scaling of diffraction intensities. Acta Crystallogr A Found Crystallogr 59, 228–234 (2003).

51. Borek, D., Cymborowski, M., Machius, M., Minor, W. & Otwinowski, Z. Diffraction data analysis in the presence of radiation damage. Acta Crystallogr D Biol Crystallogr 66, 426–436 (2010).

52. Borek, D., Dauter, Z. & Otwinowski, Z. Identification of patterns in diffraction intensities affected by radiation exposure. J Synchrotron Rad 20, 37–48 (2013).

53. McCoy, A. J. et al. Phaser crystallographic software. Journal of Applied Crystallography 40, 658 (2007).

54. Emsley, P., Lohkamp, B., Scott, W. G. & Cowtan, K. Features and development of Coot. Acta Cryst D 66, 486–501 (2010).

55. Adams, P. D., et al. *PHENIX*: a comprehensive Python-based system for macromolecular structure solution. Acta Crystallogr D Biol Crystallogr 66, 213–221 (2010).

56. Sali, A. & Blundell, T. L. Comparative protein modelling by satisfaction of spatial restraints. J Mol Biol 234, 779–815 (1993).

57. Jo, S., Kim, T., Iyer, V. G. & Im, W. CHARMM-GUI: A web-based graphical user interface for CHARMM. Journal of Computational Chemistry 29, 1859–1865 (2008).

58. Huang, J. et al. CHARMM36m: an improved force field for folded and intrinsically disordered proteins. Nat Methods 14, 71–73 (2017).

59. Lindahl Abraham, Hess & van der Spoel. GROMACS 2019.4 Manual.

60. Darden, T., York, D. & Pedersen, L. Particle mesh Ewald: An N⋅log(N) method for Ewald sums in large systems. The Journal of Chemical Physics 98, 10089–10092 (1993).

61. Hess, B., Bekker, H., Berendsen, H. J. C. & Fraaije, J. G. E. M. LINCS: A linear constraint solver for molecular simulations. Journal of Computational Chemistry 18, 1463–1472 (1997).

62. Nosé, S. A unified formulation of the constant temperature molecular dynamics methods. The Journal of Chemical Physics 81, 511–519 (1984).

63. Hoover, W. G. Canonical dynamics: Equilibrium phase-space distributions. Phys. Rev. A 31, 1695– 1697 (1985).

64. Parrinello, M. & Rahman, A. Polymorphic transitions in single crystals: A new molecular dynamics method. Journal of Applied Physics 52, 7182–7190 (1981).

65. Violin, J. D., Zhang, J., Tsien, R. Y. & Newton, A. C. A genetically encoded fluorescent reporter reveals oscillatory phosphorylation by protein kinase C. The Journal of Cell Biology 161, 899–909 (2003).

66. Akizawa, H., Lopes, E. M. & Fissore, R. A. Zn2+ is essential for Ca2+ oscillations in mouse eggs. eLife 12, RP88082 (2023).

67. Chen, V. B. et al. MolProbity: all-atom structure validation for macromolecular crystallography. Acta Crystallogr D Biol Crystallogr 66, 12–21 (2010).

