## Supplemental data for "A domain-swapped CaMKII conformation facilitates linker-mediated allosteric regulation"

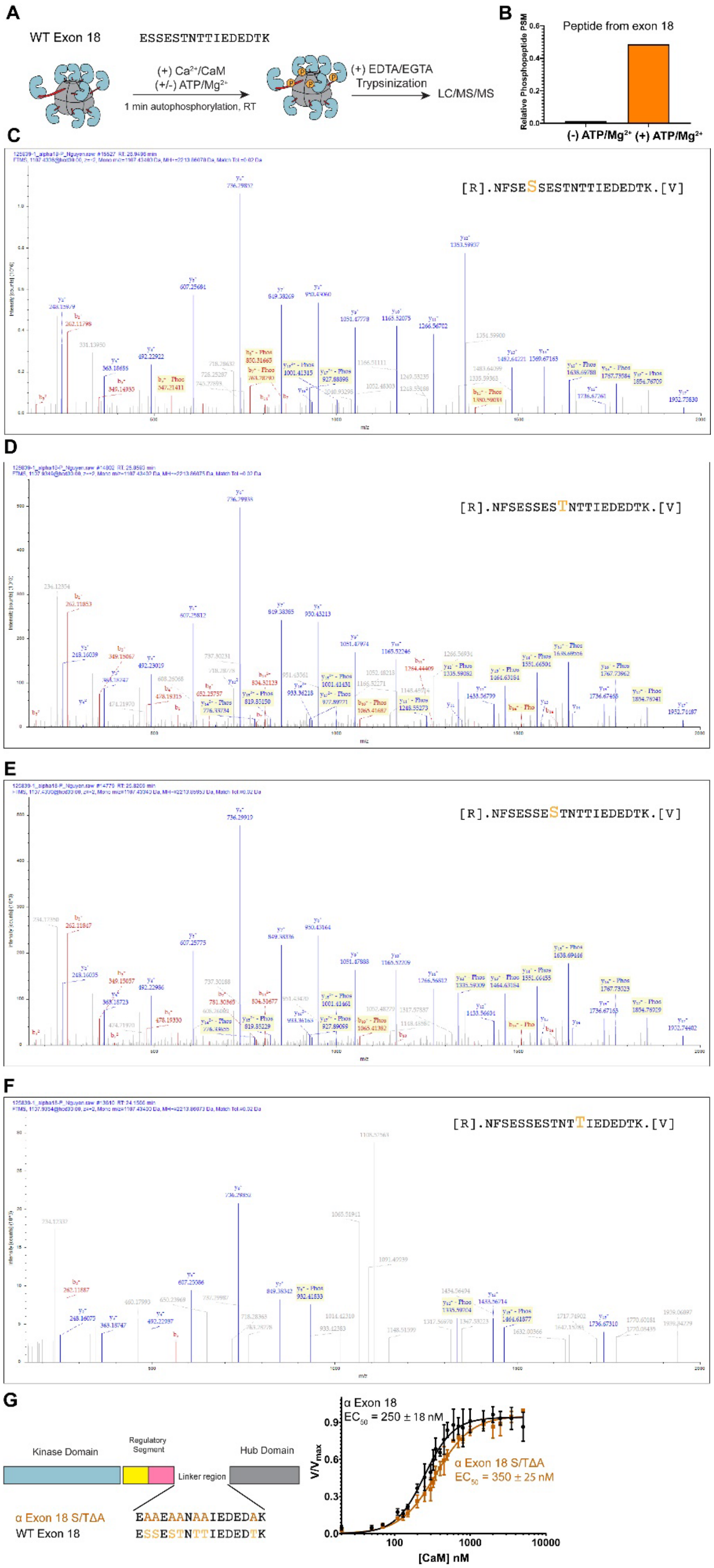
A
WT Exon 18
ESSESTNTTIEDEDTK
(+) Ca2+/CaM
(+/-) ATP/Mg2+
1 min autophosphorylation, RT
(+) EDTA/EGTA
Trypsinization
LC/MS/MS
B
Peptide from exon 18
Relative Phosphopeptide PSM
(-) ATP/Mg2+
(+) ATP/Mg2+
C
[R].NFSESSESTNTTIEDEDTK.[V]
D
[R].NFSESSESTNTTIEDEDTK.[V]
E
[R].NFSESSESTNTTIEDEDTK.[V]
F
[R].NFSESSESTNTTIEDEDTK.[V]
G
Kinase Domain
Regulatory Segment
Hub Domain
Linker region
α Exon 18 S/TΔA
WT Exon 18
EAAEAANAAIEDEDAK
ESSESTNTTIEDEDTK
α Exon 18
EC50 = 250 ± 18 nM
α Exon 18 S/TΔA
EC50 = 350 ± 25 nM
V/Vmax
[CaM] nM

**Supplemental Figure 1. Autophosphorylation of exon 18 occurs but does not change $Ca^{2+}$/CaM sensitivity.** (A) Autophosphorylation of exon 18 holoenzyme was carried out at room temperature for 1 minute in the presence of 50 nM CaM, 500 uM $Ca^{2+}$, 2 mM ATP, 10 mM $Mg^{2+}$, 25 mM Tris pH 8, 150 mM KCl. The reaction was quenched with 10 mM EGTA/EDTA and processed prior to being submitted to LC/MS/MS. (B) More peptide-spectrum matches (PSMs) of phosphorylated peptides from exon 18 were detected from samples incubated with than that without ATP/$Mg^{2+}$. Relative PSM was calculated by taking the ratio between phosphorylated peptide PSM and the non-phosphorylated peptide PSM within the same sample. (C-F) Spectra of phosphorylated peptides from exon 18. (G) Kinase assay shows that mutating all Ser and Thr of exon 18 to Ala did not shift the $EC_{50}$ to the left. Each data point of each graph represents the mean ± SD, n=3.

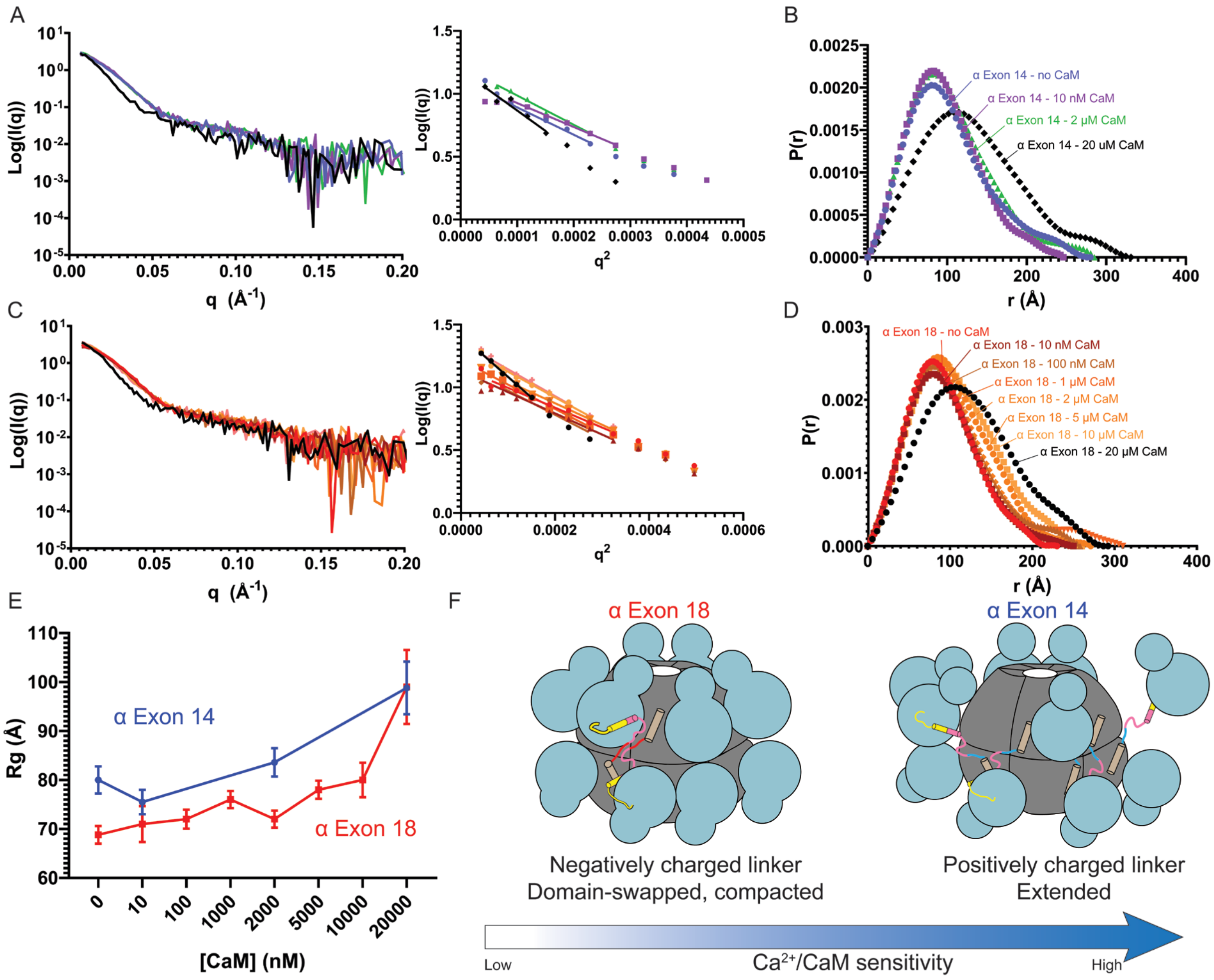


**Supplemental Figure 2. CaMKIIα exon 18 holoenzymes undergo a larger decompaction than CaMKIIα exon 14 holoenzymes upon activation.** Scattering traces of CaMKIIα exon 14 (A) and CaMKIIα exon 18 (C) are shown, and inlets are the Guinier regions used to fit the $R_g$ values. Pair distribution functions of CaMKIIα exon 14 and CaMKIIα exon 18 are shown in (B) and (D), respectively. (E) $R_g$ values of CaMKIIα exon 14 and CaMKIIα exon 18 at different concentrations of $Ca^{2+}$/CaM. Data are shown as the fitted $R_g$ ± standard deviation of the fit function (RAW v2.1.4). (F) Models showing a domain-swapped, compacted CaMKIIα exon 18 holoenzyme that is less sensitive to $Ca^{2+}$/CaM than an extended CaMKIIα exon 14 holoenzyme.

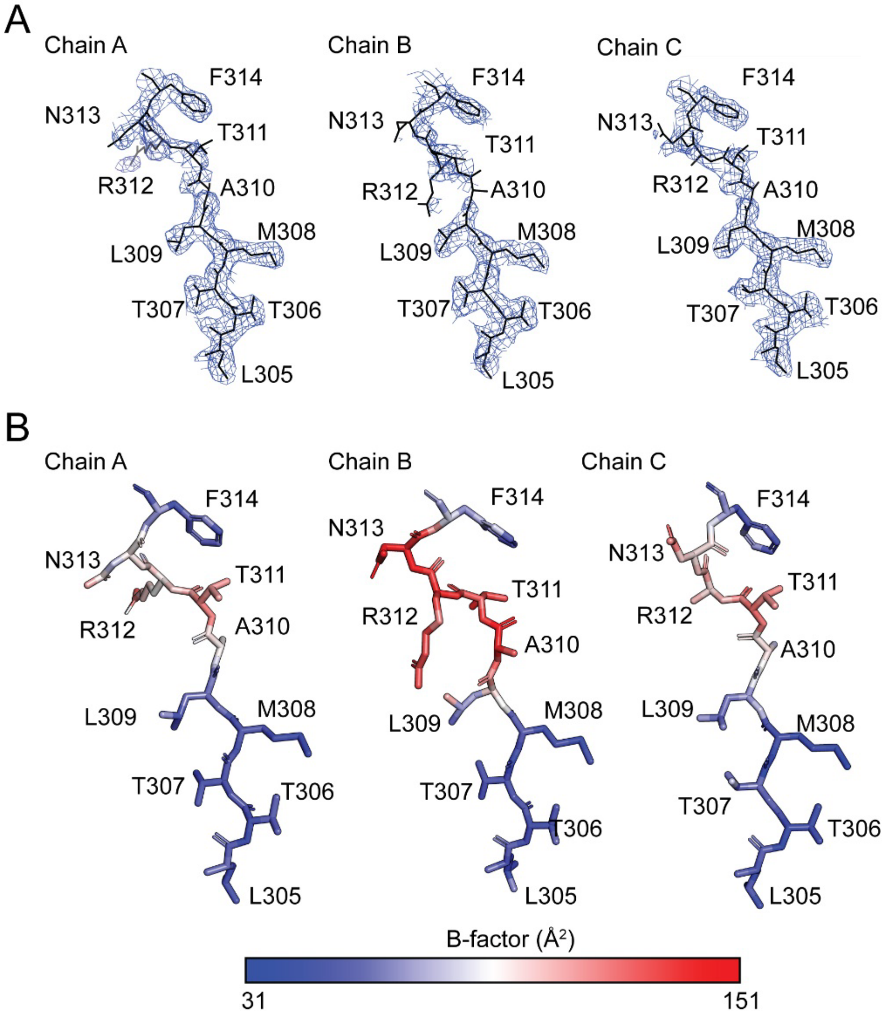


**Supplemental Figure 3. Electron density for connector segments.** (A) 2Fo-Fc map contoured at sigma 0.7 Å. For all three chains, the electron density is discontinuous for the A310 alpha carbon. (B) Chains colored by B-factor in Pymol. The density of R312 sidechain is missing.

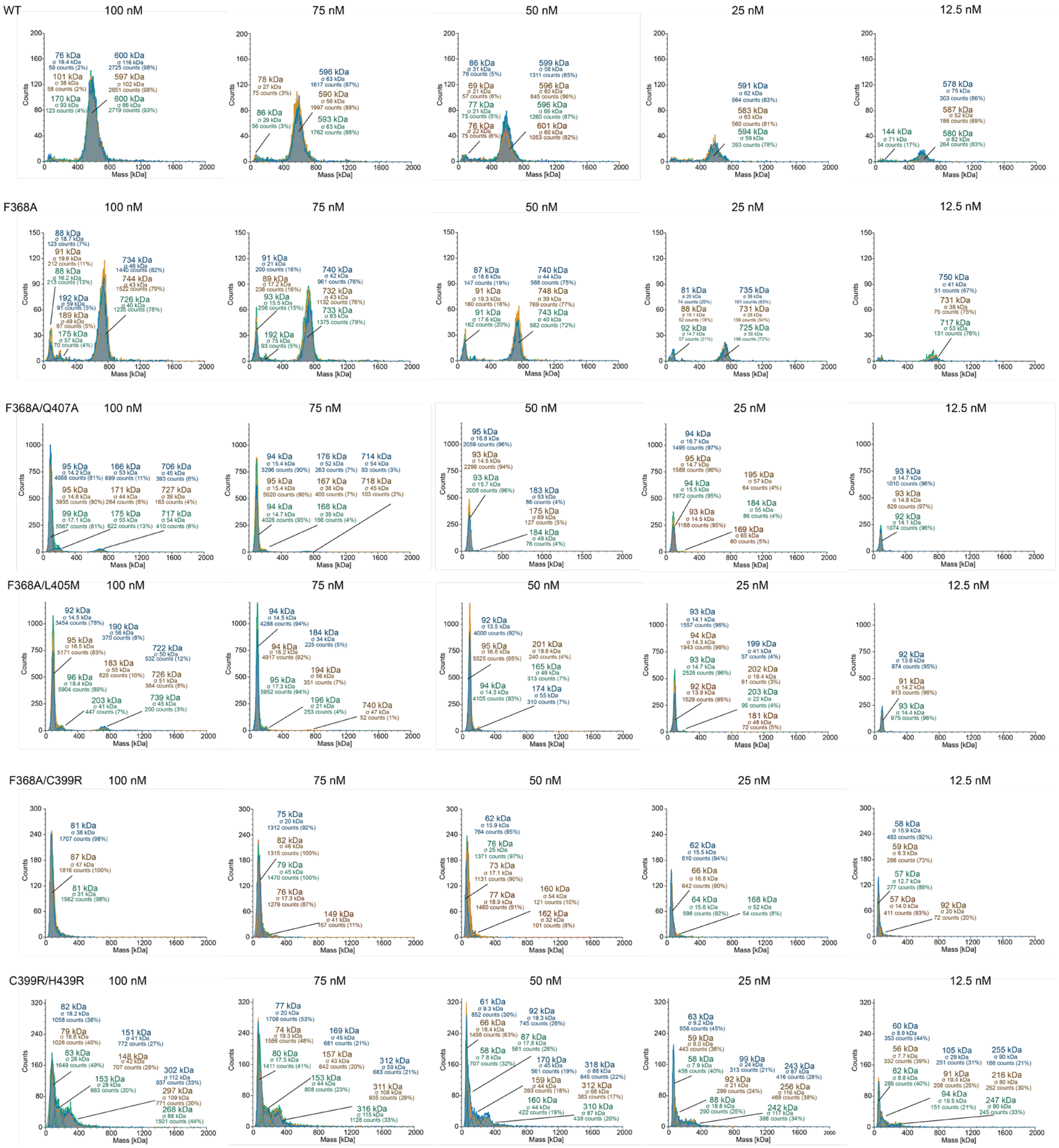


**Supplemental Figure 4.** Raw data used to generate Figure 6. Each panel contains 3 or 4 replicas for each concentration of protein.

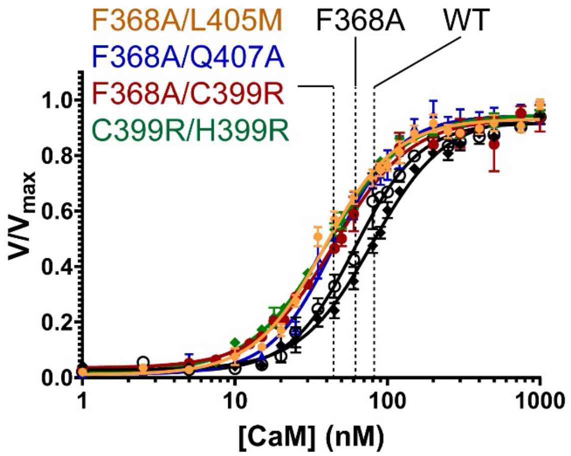


| CaMKIIδ no linker | $EC_{50}$ (nM) | h |
|---|---|---|
| WT | 80 ± 4 | 1.8 ± 0.2 |
| F368A | 61 ± 4 | 2 ± 0.2 |
| F368A/Q407A | 45 ± 3 | 2 ± 0.2 |
| F368A/L405M | 39 ± 3 | 1.8 ± 0.2 |
| F368A/C399R | 45 ± 3 | 1.7 ± 0.2 |
| C399R/H399R | 40 ± 2 | 1.7 ± 0.1 |

**Supplemental Figure 5. Activity assays of CaMKIIδ–0 wildtype and interface mutants.** Activity against syntide-2 was measured over a range of CaM concentrations for the mutants listed. Data are shown as the mean ± SD (n=3).

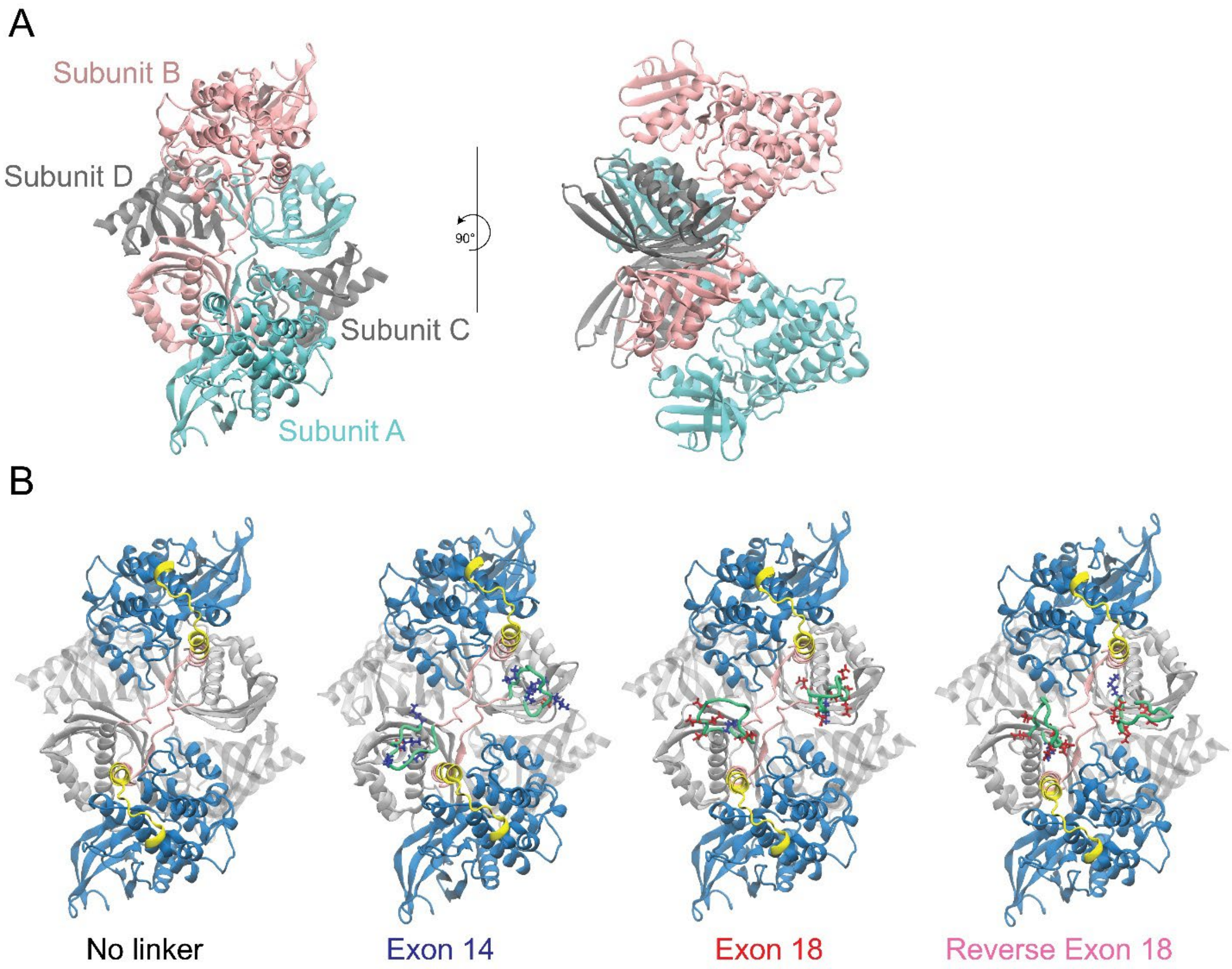


**Supplemental Figure 6. CAMKII tetramer for simulation.** (A) Tetrameric CaMKII used for linker modeling in MD simulations. This is the smallest unit which houses all possible kinase:hub interactions. Subunits A and B are full-length. Subunits C and D are hub domain one. (B) Tetrameric CaMKIIs with different linker composition. Exon 14, Exon 18, and Reverse Exon 18 were built into the no linker tetramer structure using MODELLER. Kinase: indigo; Regulatory segment: yellow and pink; Calmodulin binding region: pink; hub: gray; linkers: green. The charged residues of the linkers are represented with sticks. Positively charged residues: blue; Negatively charged residues: red.

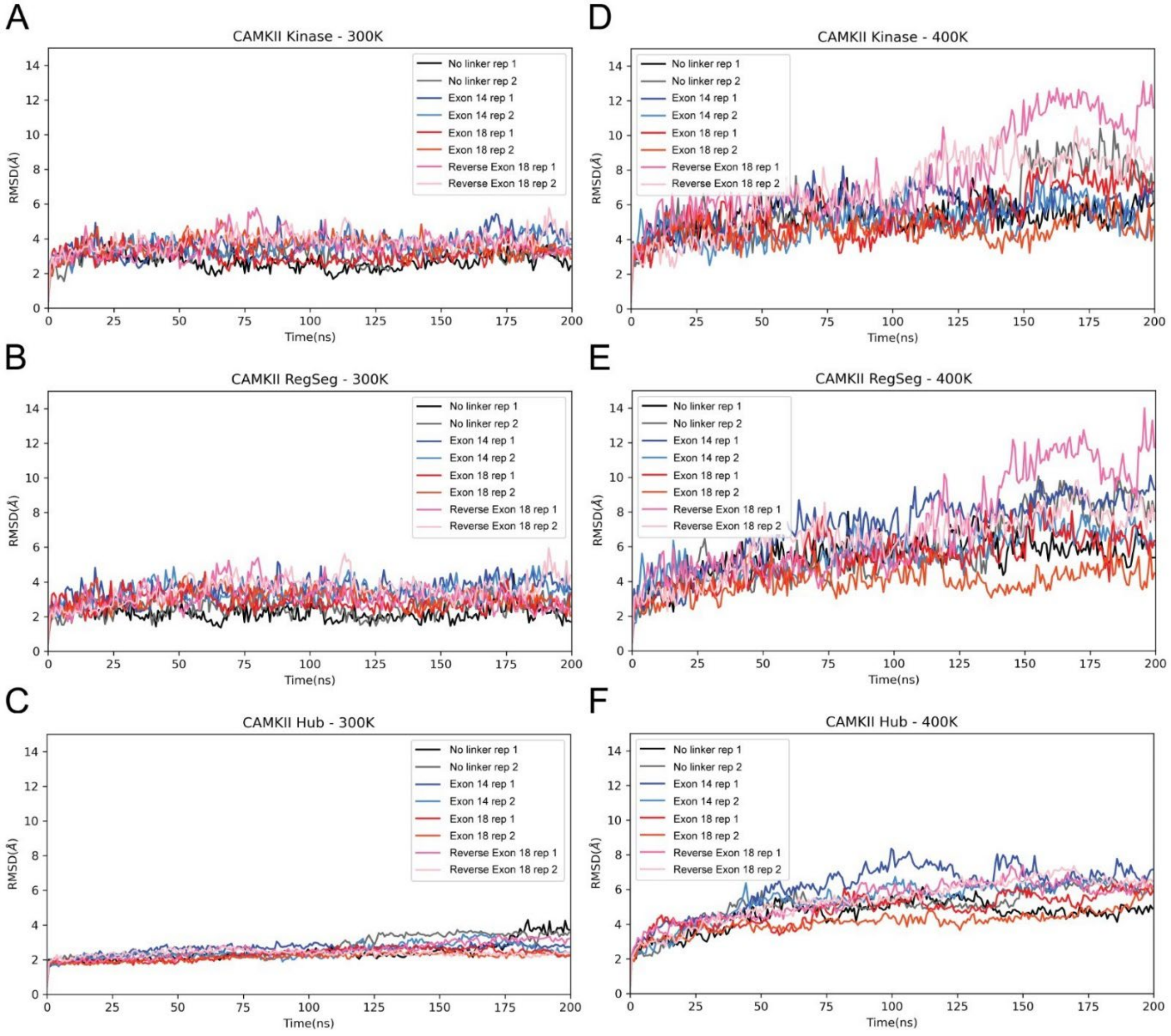


**Supplemental Figure 7. CaMKII domains are relatively stable during the 200 ns simulation.** (A-C) Root-mean-square deviation (RMSD) was calculated at 300 K for each CaMKII construct and the hub domain was used as the reference in each calculation. (D-F) Same procedure, except the temperature was set to 400 K. Each of the constructs (no linker, Exon 14, Exon 18, and Reverse Exon 18) has two replicas.

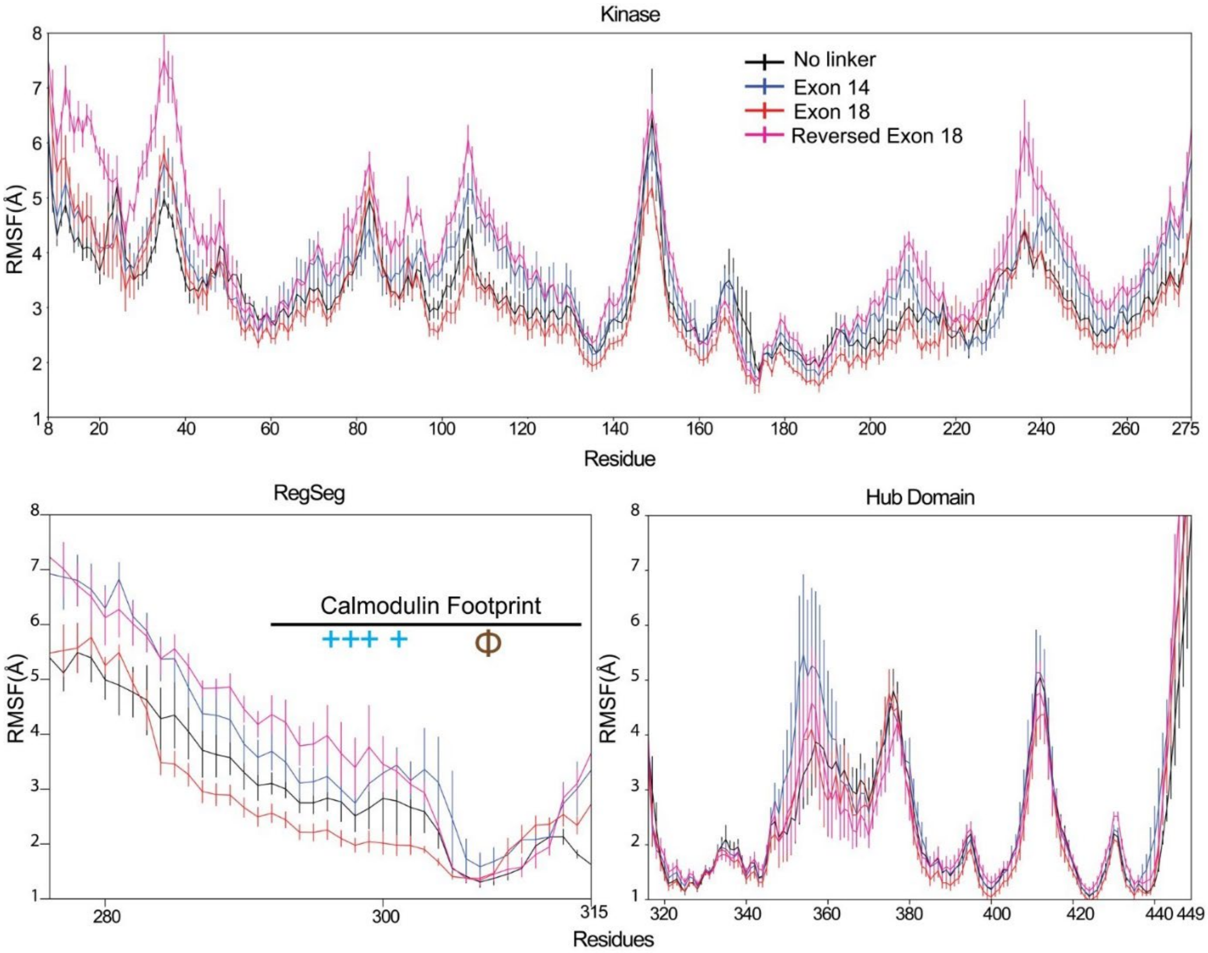


**Supplemental Figure 8**. **Linker composition alters the fluctuations of the regulatory domain.** RMSF calculation (n = 2) was carried out in the simulations of domain-swapped CaMKII models containing four subunits. On the calmodulin footprint, (+) denotes a positively charged region, and Φ denotes a hydrophobic region.

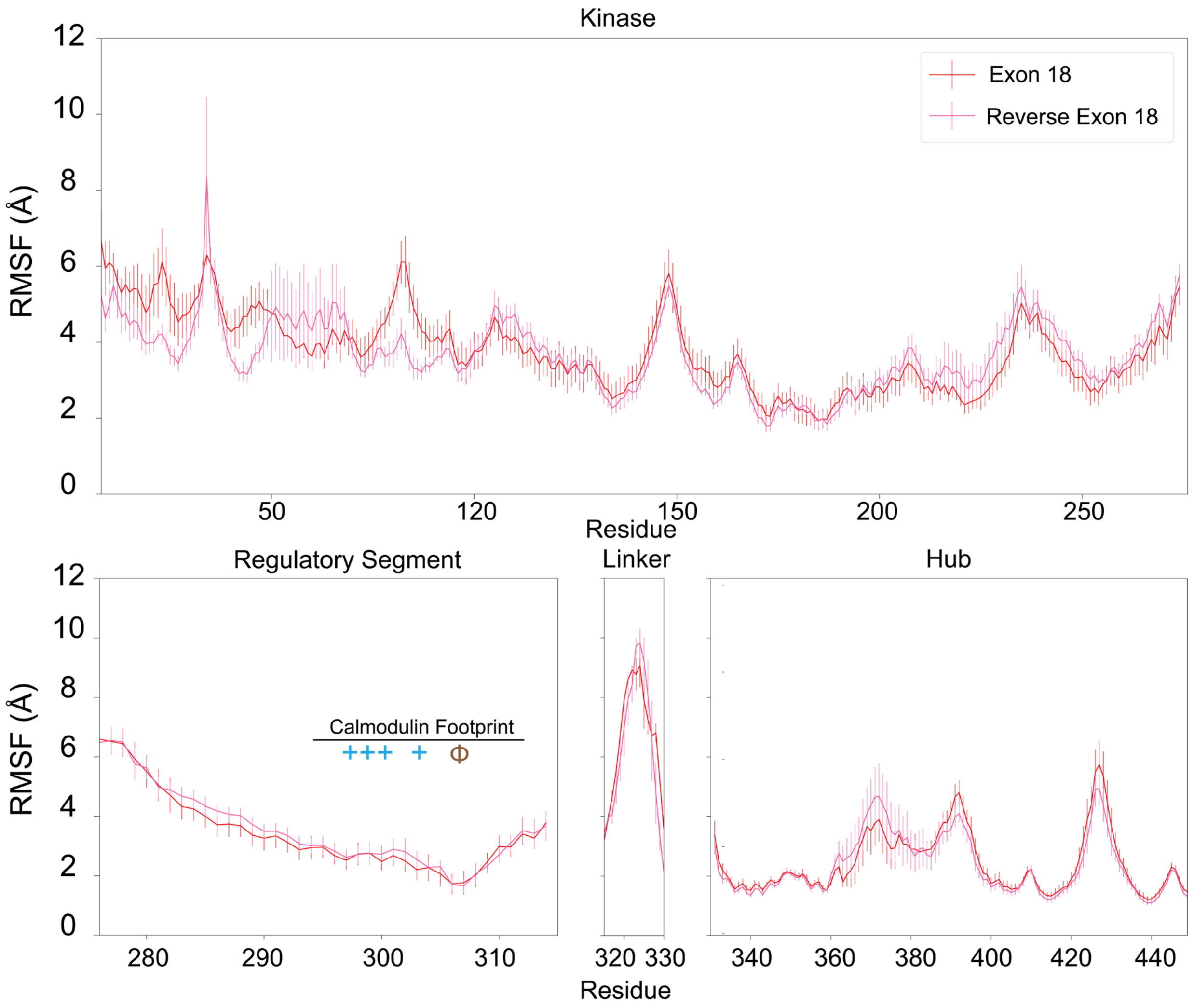


**Supplemental Figure 9**. **Domain-swapping and linker sequence facilitate regulatory segment dynamics.** In a non-swapped conformation, the differences in RMSF were not observed in between CaMKII Exon 18 and CaMKII Reverse Exon 18 (n=2). On the calmodulin footprint, (+) denotes a positively charged region, and Φ denotes a hydrophobic region.

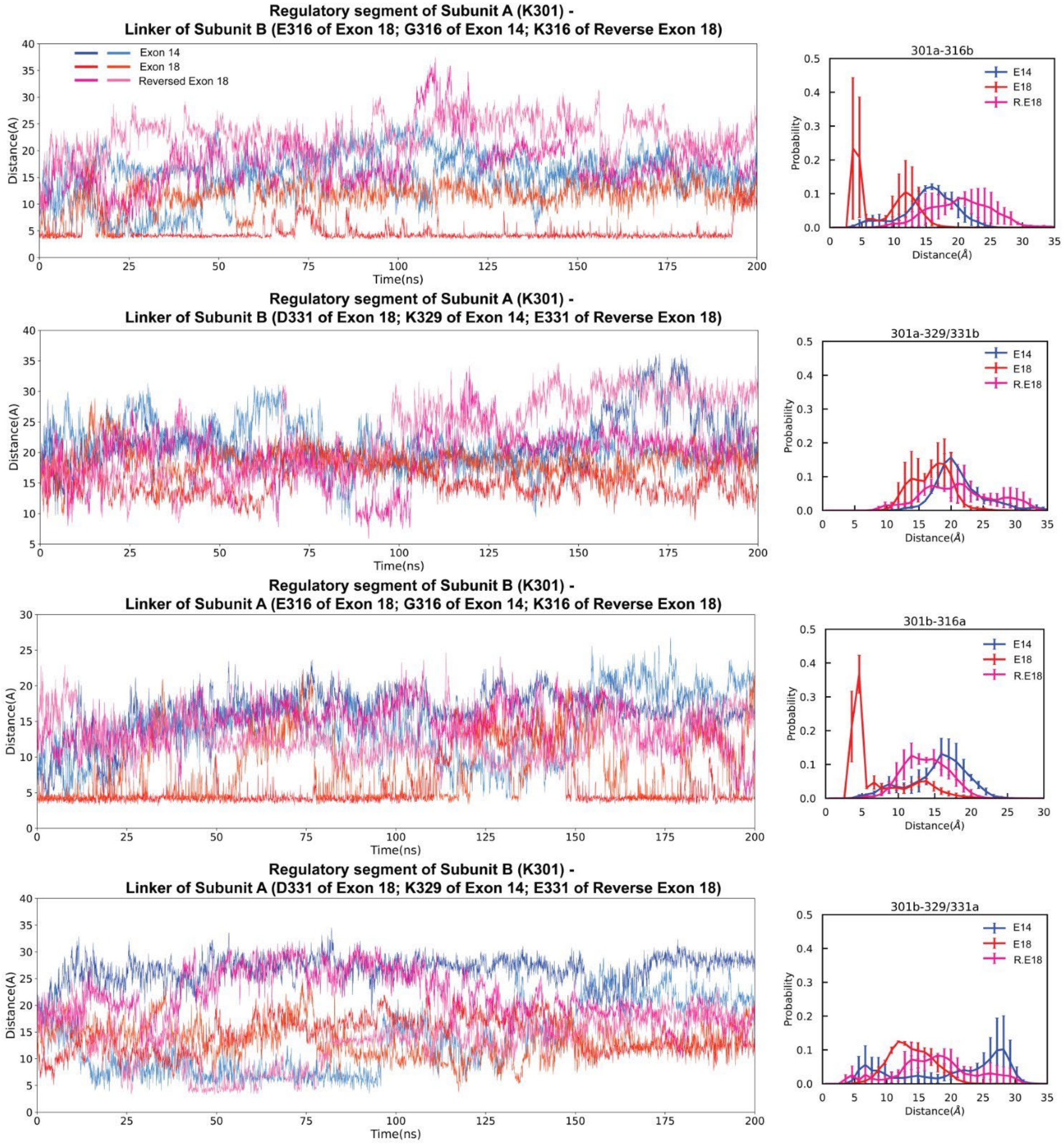


**Supplement Figure 10. There is a close interaction between the calmodulin-binding region (K301) and the N-terminal end of the variable linker.** Pair-wise interactions are plotted between K301 (calmodulin binding region) and either the N-terminal residue (residue 316) or the C-terminal residue of the linker (residue 329 for exon 14, residue 331 for exon 18 and reversed exon 18) over the 200-ns trajectory. Data from interactions between K301 of chain A and residue 316 of chain B, and vice versa, are shown. The distributions of the distance for each pair of interaction are shown in the right panel (n=2).

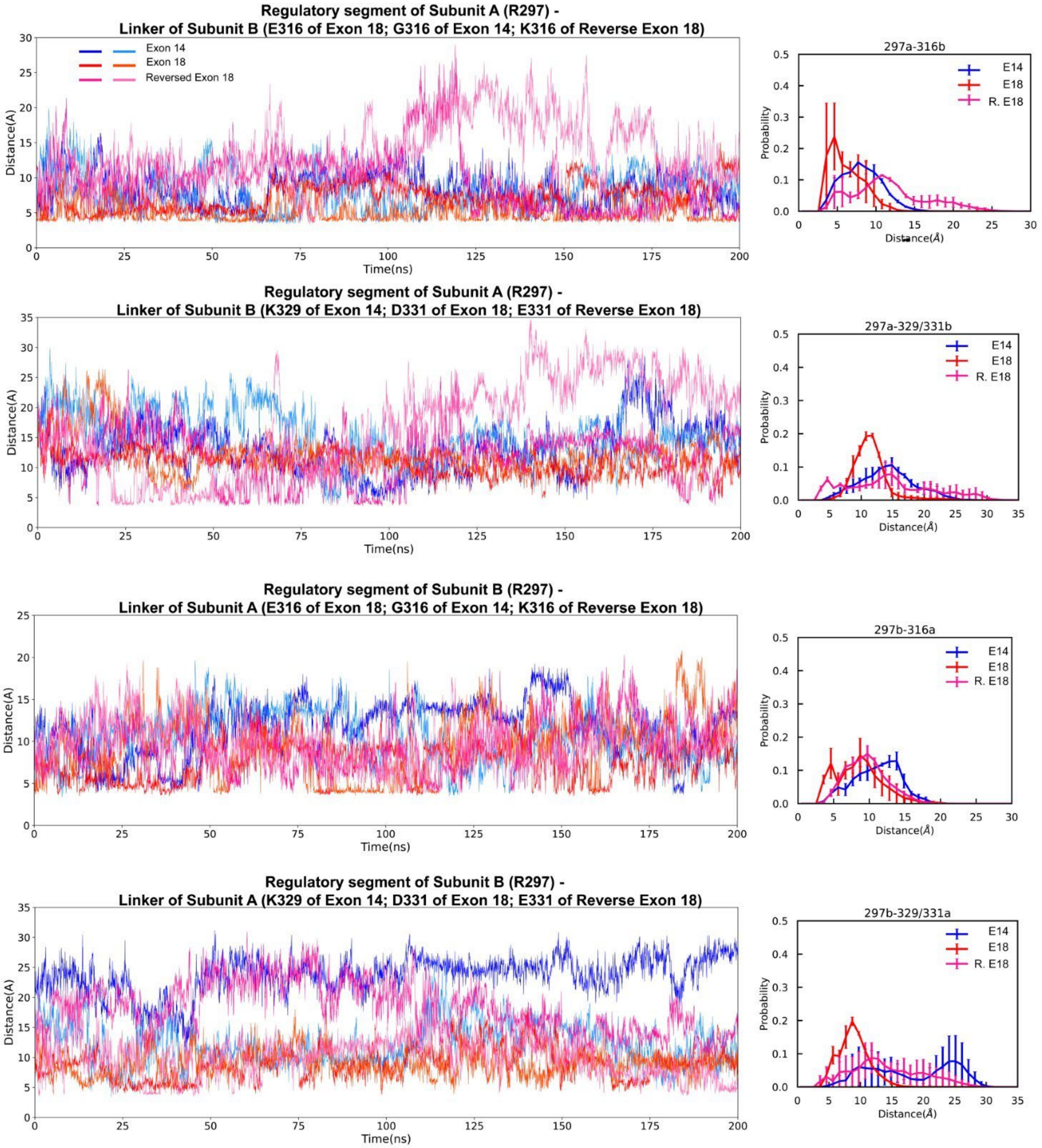


**Supplement Figure 11. There is a modest interaction between the calmodulin-binding region (R297) and the N-terminal end of the variable linker.** Pair-wise interactions plotted between R297 (calmodulin binding region) and either the N-terminal (residue 316) or C-terminal residue of the linker (residue 329 for exon 14, residue 331 for exon 18 and reversed exon 18) over the 200-ns trajectory. Data from interactions between R297 of chain A and residue 329/331 of chain B, and vice versa, are shown. The distributions of the distance for each pair of interactions are shown in the right panel (n=2).
